# ‘If the shoe fits?’ – technical nuances of plasma proteomic workflows in clinical and preclinical contexts

**DOI:** 10.64898/2026.01.12.699164

**Authors:** Samantha Emery-Corbin, Joel R Steele, Dylan H Multari, Erwin Tanuwidjaya, Iresha Hanchapola, Komagal Kannan Sivaraman, Han-Chung Lee, Scott A Blundell, Idrish Ali, Terence J O’Brien, Pouya Faridi, Ralf B Schittenhelm

**Affiliations:** Monash Proteomics & Metabolomics Platform, Department of Biochemistry and Molecular Biology, Biomedicine Discovery Institute, Monash University, Clayton, Victoria 3800, Australia; Department of Medicine, School of Clinical Science, Faculty of Medicine, Nursing & Health Science, Monash University, Clayton, VIC, Australia; Centre for Cancer Research, Hudson Institute of Medical Research, Clayton, VIC, Australia; Department of Neuroscience, School of Translational Medicine, Monash University, Melbourne, Victoria 3004, Australia

**Author notes:** Corresponding Author Information: Samantha Emery-Corbin Postal Address: Biomedicine Discovery Institute, Level 2, Building 75, 15 Innovation Walk, Monash University, Clayton, VIC 3800, Australia.

## Abstract

The push for new clinical biomarkers has seen rapid innovation in biofluid analysis, particularly for plasma. For mass-spectrometry (MS)-based analysis, achieving depth and quantitative accuracy whilst ensuring throughput continues to shape plasma methods development. Numerous workflows have emerged that mitigate high-abundance suppression and expand dynamic range, especially when paired with next-generation MS instrumentation. Yet systematic evaluations that also consider biological variables (e.g., biofluid type, species) and technical parameters (e.g., MS methods) are limited. Here, we benchmarked eight sample-preparation workflows spanning neat approaches (SP3, STrap), depletion (perchloric acid, PerCA), and corona-enrichment strategies (MagNet HILIC/SAX, Enrich-iST, ProteoNano). We compared their performance across human plasma, human serum, and rat plasma, analysing all samples on an Orbitrap Astral (Thermo) using two plasma-optimised data-independent acquisition (DIA) methods: one discovery-maximised and one throughput-maximised. We identified 2,726 human and 3,767 rat proteins across workflows and methods, including ∼1,000 from neat plasma. Increasing throughput incurred a ∼20-30% reduction in depth, depending on workflow and species. EV-enrichment produced the deepest proteomes but with distinct compositions relative to neat, depleted, and secreted-protein-enriched samples, revealing a unique sub-proteome niche. Several workflows also performed markedly better in rat plasma, supporting improved sensitivity for preclinical analyses. Enrichment or depletion dramatically reshaped the balance of tissue-and cell-specific proteins detectable in plasma, suggesting that workflow choice should be guided by the organs, immune targets, or inflammatory signals most relevant to the study. In this vein, statistical analysis of differentially abundant proteins showed that >90% of detected proteins were significantly altered between workflows, with the largest numbers arising from the corona-enrichment strategies, underscoring how strongly workflow choice shapes the downstream proteome. Taken together, these findings emphasise a rapidly expanding plasma methodological landscape, where the most effective workflow is the one most precisely tailored to a cohort’s biology.

## Introduction

Biomarkers are molecular indicators that span the clinical continuum between healthy and disease states. Among these, protein biomarkers claim the largest share of molecules assayed for clinical diagnostics, with plasma being the predominant source of patient-derived samples (1). Sampling plasma is routine, inexpensive and minimally invasive, supported by robust and well-established collection protocols (2–4). Arguably, the current suite of FDA-approved plasma biomarkers represents only a fraction of plasma’s diagnostic potential, given plasma’s rich compositional proteome from human tissue and immune cell sources (5). This is underscored by recent studies reporting quantitative detection of over 5,000 and even 10,000 proteins in plasma (6, 7).

For proteins specifically, the disconnect between the number of potential and translated plasma biomarkers is attributed to its challenging dynamic range (8), particularly in a field historically driven by mass spectrometry (MS) based discovery (9). The canonical or core plasma proteome consists of several hundred abundant proteins (10) that are readily detected despite the suppressive effects of the top 20 proteins, which together account for 99% of plasma by mass. Consequently, the aspirational peak of plasma proteomics (or inversely, the iceberg’s tip) is the other 1% of mass which contains tens of thousands of lower-abundance proteins, including those derived from tissue leakage and immune responses, which may hold greater diagnostic value across the clinical continuum.

Plasma proteomics has seen incredible innovation in recent years. Between 2021 (10) and 2024 (11), the HUPO Plasma Proteome Project (HPPP) reference PeptideAtlas build grew its peptide-spectrum matches (PSMs) and distinct peptides by 83% and 27%, respectively, and its canonical proteins increased 20-fold from 213 to 4,608. This growth coincided with major advances in MS technology, including the shift from data-dependent to data-independent acquisition (DDA to DIA) (12, 13), and the introduction of new MS analysers (14, 15). Engineered functionalised nanoparticles (NPs) (16, 17) for plasma dominated new sample enrichment methods, firstly the Proteograph XT (Seer) (18), followed by MagNet (ReSyn Biosciences) (19) and ENRICH (PreOmics) (20), and more recently ProteoNano (Nanomics) (21). Although ten years ago over half of the plasma studies used top-10/20 depletion columns (1), modern depletion favours selective precipitation methods such as perchloric acid (PCA) (22, 23). The convergence of these innovations – spanning MS technologies, workflows and automation – has enabled large-scale plasma proteomics studies to expand dramatically, with cohort sizes growing from just over 1,000 (24) to 36,000 (23) plasma samples in the last five years. These studies now routinely achieve proteome depths that far exceed previous benchmarks.

Given MS-based proteomics concatenates independent modules spanning sample preparation, liquid chromatography (LC) and MS methods, the scope of a current plasma workflow is substantial and highly multivariate. Unsurprisingly, numerous method-benchmarking studies have arisen in the scholarship (7, 20, 25–27) and pre-scholarship (28, 29). Collectively, these studies are building comprehensive references across neat, enriched, and depleted plasma approaches, and provide key insights related to their adaptability, applicability, and suitability for specific use cases. More specifically, some have also evaluated technical and sample collection biases (27, 30), collection tube (27, 29), and proteome depth across sample per day (SPD) cut-offs (28). Such benchmarking is essential for assessing the practical applicability of workflows across different laboratories, and will have ripple effects across future protein biomarker discovery (7).

In this study, we contribute to this growing body of knowledge by benchmarking sample workflows across both human and rodent biofluids. We evaluated eight sample preparation methods using human serum and plasma, alongside rodent samples (rat plasma), highlighting their performance in preclinical models. Samples were analysed on the Orbitrap Astral mass spectrometer (Thermo Scientific) using two methods with optimised LC and MS settings: a longer ‘Discovery’ method (16 SPD, including washes and QC) and a ‘Throughput’ method (32 SPD, including washes and QC). Our findings demonstrate that sample and instrument workflows profoundly shape the plasma proteome—within species, between species, and across biofluids – where there’s no one-size-fits-all solution, and workflows must be tailored on a case-by-case basis.

## Experimental Procedures

### Experimental Setup

Blood was obtained from healthy donors, following approval of the Monash University Human Research Ethics Committee (HREC number = 13019A). For Rats, baseline blood was collected from 11-week-old Sprague–Dawley rats obtained via the Monash Animal Research Platform as part of the EpiBios4Rx project. All procedures were approved by the Alfred Medical Research & Education Precinct Animal Ethics Committee (E/1799/2018/M). Animals were anesthetised with isoflurane (5% induction, 2% maintenance) as previously described (31), and blood collected from the lateral tail vein using a 23G butterfly needle into K2EDTA tubes. Samples were centrifuged at 1,300 × g for 10 min at 4 °C to isolate plasma, which was aliquoted (50 µL), flash-frozen on dry ice, and stored at −80 °C until analysis.

### Neat Plasma Workflows

Neat plasma was prepared using the STrap (32) and SP3 protocols (33). For both protocols, 2 µL of neat plasma was solubilised in 100 µL of lysis buffer of 5% SDS lysis buffer containing 10 mM Tris (2-Carboxyethyl) phosphine (TCEP) and 40 mM 2-chloroacetamide (2-CAA). After protein reduction/alkylation was performed for 10 minutes (min) at 95 °C, a 25 µL aliquot was taken from the lysed solution for the subsequent digestion steps (∼35 µg).Briefly, for STrap proteins were acidified with 2.5 µL of 12% phosphoric acid (final ∼1.2%), then 175 µL of binding buffer (90% methanol in 50 mM TEAB; pH 7.1) was added and mixed thoroughly. The mixture was loaded onto STrap micro spin columns centrifuged at 4,000 × g for 30 sec, discarding the flow-through. Columns were washed three times with 400 µL binding buffer (4,000 × g, 30 sec each). For overnight digestion, Trypsin Gold (MS-grade, Promega) and Lys-C (FUJIFILM Wako) was prepared at 0.5 µg/µL in 50 mM TEAB and 1 µg total was added to each column. Digestion proceeded at 37 °C overnight without shaking. Peptides were eluted sequentially with 80 µL 50 mM TEAB, 80 µL 0.2% Formic acid (FA) in water, and 80 µL 50% acetonitrile (ACN) with 0.2% FA, and the eluates were dried down via vacuum centrifugation in a CentriVap (Labconco) prior to SPE clean-up.

For SP3, proteins were subjected to on-bead precipitation with 20 µL of Sera-Mag™ SpeedBeads (Cytiva) by adding acetonitrile (ACN) to a final 70% (v/v) and mixing for 20 min at room temperature on a ThermoMixer C (450 rpm). Samples were placed on a magnetic rack, supernatants removed, and beads washed three times (200 µL each): twice with 70% ethanol and once with 100% ACN. Residual solvent was evaporated via vacuum centrifugation in a CentriVap (Labconco), after which digestion buffer (100 mM Tris-HCl, pH 8.0) containing Lys-C (FUJIFILM Wako) and SOLu-Trypsin (MS-grade, Promega), each at 0.5 µg/µL in 50 mM TEAB concentration was added using 1 µg total for each. On-bead digestion was carried out for 1.5 h at 37 °C with shaking at 450 rpm, after which digests were transferred off beads and acidified by adding 10% tri-fluoro acetic acid (TFA) to a final concentration of 1%.

### MagNet Workflows

MagNet bead enrichment (34) was performed using 12.5 µL of either MagNet HILIC, MagNet SAX or a 50:50 combination of HILIC/SAX beads (ReSyn BioSciences). For each sample, 50 µL of spun, neat plasma were combined with 50 µL equilibration buffer (100 mM Bis-Tris-propane, pH 6.3, 150 mM NaCl) and 12.5 µL of pre-washed MagNet beads (wash buffer: 50 mM Bis-Tris-propane, pH 6.3, 150 mM NaCl). Corona enrichment was performed at room temperature for 30 min on a ThermoMixer C (300 rpm), placed on a magnetic rack for 1-2 min to capture beads, and washed three times with 500 µL wash buffer (5 min mixing at 300 rpm per wash). After completion of washes, 50 µL lysis buffer (2% SDS, 100 mM Tris-HCl pH 8.5, 10 mM TCEP, 40 mM 2-CAA) were added to the bead pellet and reduction/alkylation proceeded at 40 °C for 45 min with gentle shaking (300 rpm). Protein aggregation capture was then initiated by adding ACN to 70% (v/v; 130 µL ACN) and incubating 20 min at room temperature (300 rpm). Beads were re-captured on a magnet (1–2 min) and washed three times (200 µL each): twice with 70% ethanol and once with 100% ACN, removing residual solvent after each wash. Beads were dried (brief lyophilization or 5–10 min air-dry), then 50 µL digestion buffer (100 mM Tris-HCl, pH 8.0) containing Lys-C (FUJIFILM Wako) and SOLu-Trypsin (MS-grade, Promega), each using 1 µg at 0.5 µg/µL concentration were added. On-bead digestion proceeded for 1.5 h at 37 °C with shaking at 300 rpm, after which digests were transferred off beads and acidified by adding 10% TFA to a final concentration of 1%.

### EnrichIST Workflow

EnrichIST (PreOMICS) was performed according to manufacturer instructions. Briefly, 20 µL of spun plasma were processed. EN-BEADS (25 µL) were washed three times with EN-WASH, combined with 80 µL EN-BIND and 20 µL plasma, and incubated for 30 min at 30 °C on a Thermomixer C (1,200 rpm), followed by three EN-BIND washes. Beads were lysed with 50 µL LYSE-BCT (95 °C, 10 min), cooled, and DIGEST freshly reconstituted in RESUSPEND-BCT was added (50 µL per reaction) for on-bead digestion (37 °C, 90 mins, 1,200 rpm). Reactions were stopped with 100 µL STOP, peptides were purified on the supplied cartridges, dried, and stored at −80 °C prior to reconstitution.

### ProteoNano Workflow

The ProteoNano protocol was followed according to manufacturer’s instructions, as adapted for manual processing. A total of 40 µL plasma was transferred to a low-bind tube and mixed 1:1 with Buffer B (150 mM KCl, 0.05% CHAPS, Tris-EDTA pH 7.4). Reagent A (magnetic nanobeads) was equilibrated at room temperature and vortexed to homogeneity. For manual processing, for each sample 100 µL beads were transferred to a fresh tube, magnetically settled, and 80 µL supernatant removed to yield a 20 µL concentrated bead suspension. The 20 µL Reagent A were added to the 80 µL sample mixture (final composition: 40 µL sample: 40 µL Buffer B: 20 µL beads), then incubated 1 h at room temperature with shaking (1,200 rpm). Samples were placed on a magnetic rack for 1-2 min to capture beads, and washed three times with 180 µL Buffer B, fully removing the final wash. For denaturation/reduction, 20 µL of Digestion Solution 1 (5 mM DTT in 50 mM ammonium bicarbonate) were added and samples were shaken at 55 °C, 1,200 rpm for 30 min. After a brief spin and cooling to room temperature, 20 µL of Digestion Solution 2 (20 mM iodoacetamide and 50 ng/µL trypsin in 50 mM ammonium bicarbonate) were added and on-bead proteolysis proceeded at 37 °C (1,200 rpm) for 3 h in the dark, after which digests were transferred off beads and acidified by adding 10% TFA to a final concentration of 1%.

### Perchloric Acid precipitation Workflow

PerCa as performed as according as previously described (35), with some amendments to accommodate a lack of in-line sample clean-up. 50 µL of spun plasma were diluted with 450 µL water in a low-bind 1.5 mL tube. Perchloric acid was added to a final concentration of 4% (v/v) with vigorous mixing to precipitate proteins, and samples were incubated at −20 °C for 15 min. Precipitates were pelleted by centrifugation (3,200 × g, 60 min, 4 °C). Carefully avoiding the pellet, 400–450 µL supernatant were transferred to a fresh tube and acidified with 40 µL 1% TFA.

Acidified supernatants were desalted on Oasis HLB SPE cartridges (10 mg sorbent; Waters) conditioned with 300 µL methanol and equilibrated with 0.1% TFA (2 × 500 µL). Samples were loaded, washed twice with 500 µL 0.1% TFA, and eluted with 100 µL 90% ACN, 0.1% TFA. Eluates were frozen and dried to completeness in a CentriVap (Labconco) vacuum concentrator, then taken forward to the STrap digestion workflow by reconstituting in the first buffer as per the previous section.

### LC-MS/MS

For samples from all workflows excluding EnrichIST, samples underwent SPE clean-up using in-house SDB-RPS (Empore) as previously desribed (36). Subsequently, all samples were reconstituted in MS-ready buffer containing iRT peptides (Biognosys) at 1:600 ratio. All samples were nano-dropped (A205 Scope, NanoDrop One, Thermo Fisher Scientific) to adjust injection volumes for 300 µg and 150 µg peptide mass on column for the Discovery and Throughput LC-MS methods, respectively. All raw files and their respective Spectronaut search outputs can be accessed via PRIDE with the identifier PXD072131.

For the ‘Discovery’ method, peptides were loaded by direct injection onto Aurora® Ultimate™ 25×75 C18 UHPLC column (IonOpticks) maintained at 50 °C. Mobile phase A consisted of water with 0.1% FA, and mobile phase B was 80% ACN with 0.1% FA. Chromatographic separation was achieved over a 65-min gradient at 300–700 nL/min, beginning at 4% B, increasing to 7% B at 2.1 min, 32% B at 52.2 min, 50% B at 55.7 min, and 99% B for column washing (59.7–60 min), followed by re-equilibration to 65 min. Samples were injected using direct-injection mode with the autosampler maintained at 7 °C. The Orbitrap Astral was operated in positive-ion nano-ESI mode with static spray voltage (2000 V) and an ion transfer tube temperature of 300 °C. MS1 spectra were acquired in the Orbitrap at 240,000 resolution over m/z 380–1000 with a 10 ms transient, using profile data and a normalized AGC target of 500%. DIA spectra were acquired in the Astral analyser using a stepped, user-defined isolation window scheme (custom variable windows), with a normalized HCD collision energy of 25% and a scan range of m/z 150–1500. DIA transients were acquired at 6.1 ms with centroid data, normalized AGC target of 800%, and 0.6 s cycle time. Advanced peak determination was enabled, and all data were acquired in peptide mode with standard pressure settings.

For the ‘Throughput’ method, peptides were loaded by direct injection onto Aurora® Elite™ 15×75 C18 UHPLC column (IonOpticks) maintained at 50 °C. Mobile phase A consisted of water with 0.1% FA, and mobile phase B was 80% ACN with 0.1% FA. Chromatographic separation was achieved over a 30-min gradient at a flow rate of 400–700 nL/min, beginning at 4% B (0–6 min), increasing to 34% B at 21.1 min, then to 80% B at 24.2 min, followed by a column wash at 99% B (26.3–28 min) and re-equilibration to 30 min. The Astral was operated in positive-ion nano-ESI mode with a spray voltage of 1900 V and ion transfer tube temperature of 300 °C. MS1 spectra were acquired in the Orbitrap at 240,000 resolution over m/z 380–980 with a 20 ms transient, using profile mode and a custom AGC target (normalized AGC 500%. DIA was performed using 199 variable windows (3 m/z isolation width), acquired in the Astral analyser at 3 ms transient and 0.6 s cycle time. DIA scans used centroid data, normalized HCD collision energy of 25%, and a precursor mass range of m/z 380–980 with fragment ion scanning from m/z 145–1450. Advanced peak determination was enabled. All data were acquired in peptide mode with standard pressure settings.

### Database searching and data Analysis

The proteomic datasets were searched using a DIA-library free approach using Spectronaut (version 19.4.241120). Data files were searched against the UniProt human proteome (UP000005640, reviewed sequences, downloaded January 2025) and Rat proteome (UP000002494, one protein per gene, downloaded January 2025) using a protease specificity of Trypsin allowing oxidation of methionine as a variable modification and carbamidomethylation set as a fixed modification of cysteine. Quantitation used MaxLFQ and the method was BGS standard, sparse, no imputation, cross-run normalisation enabled. Q-value was set to 1% and protein false discovery rate (FDR) cutoff was 1%. A DIA-Analyst compatible protein group matrix was downloaded for further pre-processing as outlined below.

Enrichment of tissue-specific proteins in each workflow was assessed by one-sided hypergeometric tests against the global tissue atlas background, implemented in R via phyper. Raw p-values were corrected per workflow using the Benjamini–Hochberg FDR correction (p.adj < 0.05). Marker-specific UniProt identifications were first classified as “cell” or “tissue” based on a curated plasma atlas and for each method, the summed intensity of cell- and tissue-specific markers was compared to the corresponding global background fraction via one-sided binomial tests, with p-values adjusted by the Benjamini–Hochberg FDR correction. To perform this for the rat proteins, we first mapped rat UniProt entries to its human counterparts via three orthology resources (Ensembl GeneTrees, OMA, and HOGENOM). We also extracted the corresponding human UniProt IDs and their tissue- or cell-specific labels (5). For each rat protein, we assigned a Consensus_Label (the tissue/cell annotation agreed upon by at least two databases, or ‘FLAG’ when no agreement was reached) and a Consensus_UniProt ID (the human UniProt accession matching that consensus); proteins lacking any orthologous hit remained unannotated.

For statistical analysis of differentially abundant proteins (DAPs), raw protein intensities were log_2_-transformed and within-group cyclic-loess normalisation was applied using the normalizeCyclicLoess function (method = “fast”) with a span parameter of 0.7 and three iterations. Within each condition group, missing values were then imputed using the MinProb algorithm (impute.MinProb) with q = 0.01, replacing missing entries by draws from the lower-tail (1st percentile) distribution of that group’s observed intensities. This matrix was then analysed via the Analyst Suites (37) using DIA Analyst v0.10.4 (https://analyst-suites.org/apps/dia-analyst/) with no normalisation and no imputation selected. Functional enrichment was performed using STRING-DB (38). These data frames were also uploaded to the Baize: Plasma and Serum Contamination Assessment Tool (v3.0) (17) for evaluation of platelet, erythrocyte and coagulation contamination between workflows (https://www.guomics.com/software/Baize ).

### Experimental Design and Statistical Rationale

All experiments were performed using three independent biological replicates of pooled human plasma, pooled human serum, and pooled rat plasma (n = 3 per matrix). Pools were generated from healthy donors or control animals to provide a representative basal plasma proteome and to minimise low-abundance, donor-specific variability. After pooling, samples were aliquoted and processed independently to generate technical replication. All workflows were processed on the same day to maintain identical freeze-thaw histories and stability profiles. Sample preparation (including SDB-RPS SPE) and LC-MS analysis were batched where possible; MS injections were randomised, wash runs were inserted between samples, and instrument QC standards (HeLa digest, Pierce) were acquired every 24 hours to monitor stability. For quantitative analysis, cross-run normalised values from Spectronaut were used. To stabilise replicate means within each workflow prior to statistical analysis, we applied within-group cyclic loess normalisation and within-group MinProb imputation, avoiding across-workflow overnormalisation to preserve true method-specific differences. Statistical testing was performed in DIA Analyst using a two-sided t-test with Benjamini-Hochberg FDR correction, and significance was defined as adjusted p < 0.05 with log₂ fold change ≥ 1.

## Results

### Evaluation of workflows, logistics and peptide retrieval

We compared eight plasma workflows in clinical biofluids (human serum and EDTA plasma), and a preclinical model (rat plasma) (Figure 1A). Technical triplicates were pooled from healthy donors for required volumes, and three technicians manually prepared all samples in parallel across two days (n = 72). Workflows included two neat (STrap, SP3), five corona enrichments (MagNet SAX, MagNet HILIX, MagNet SAX+HILIC combination, EnrichIST, ProteoNano) and one depletion method (PerCa) (Supplementary Table 1). We note the ProteoNano workflow is usually automated, and designed for human plasma only.

**Figure 1.**
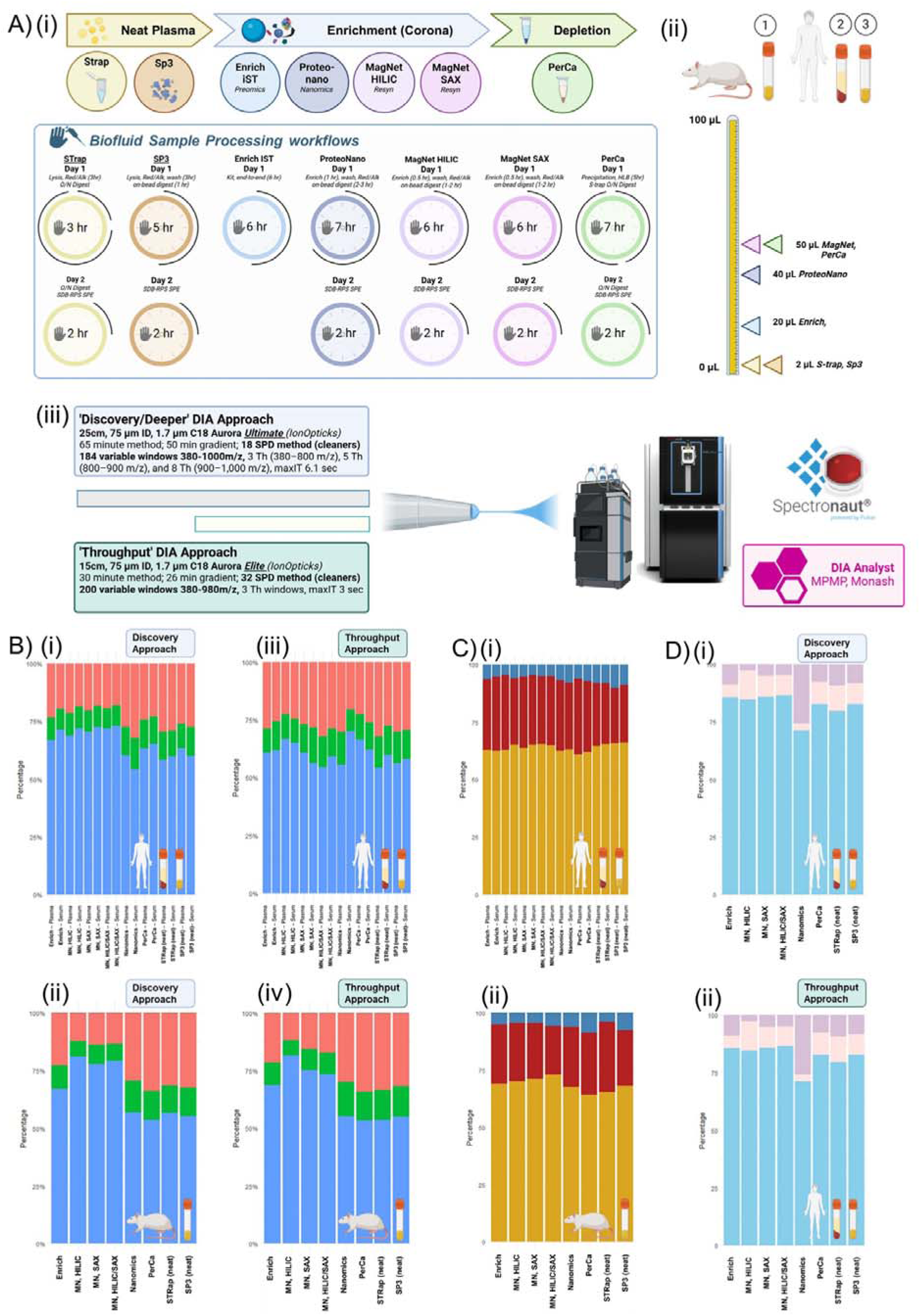
**Overview of plasma proteomic workflows and identification metrics.** (A) Diagram of sample-processing workflows and DIA LC–MS methods. Figure created with BioRender.com. (i) Eight workflows: two digestions of neat plasma (STrap, SP3), five corona-based enrichments (EnrichIST, ProteoNano, MagNet-HILIC, MagNet-SAX, MagNet-HILIC/SAX), and one depletion (PerCa). Hands-on time is shown (overnight digestions excluded). (ii) Biofluid inputs per workflow: rat plasma, human serum, and human plasma. (iii) Two plasma-optimised DIA methods: a discovery-maximised method (65 min; “Aurora Ultimate”, IonOpticks) and a throughput-maximised method (30 min; “Aurora Elite”, IonOpticks). (B) Stacked bar charts of protein identifications by workflow for: (i) human biofluids—Discovery method; (ii) human biofluids—Throughput method; (iii) rat plasma—Discovery method; (iv) rat plasma—Throughput method. (C) Stacked bars showing unique vs shared protein identifications between Discovery and Throughput methods for (i) human biofluids and (ii) rat plasma. (D) Stacked bars partitioning human biofluid proteins into shared, plasma-unique, and serum-unique sets for (i) the Discovery method and (ii) the Throughput method.

EnrichIST, being kit-based, was completed within a single day including in-kit solid-phase extraction (SPE). Single-day processing was also feasible for workflows using on-bead digestion (corona enrichments and SP3), although SPE was deferred to day 2 to avoid batch effects in workflows requiring overnight digest (Strap, PerCa). All workflows involving magnetic beads could be expedited through automated liquid-handler systems (28). Additional steps in the PerCa workflow, such as HLB clean-up (35), made it the most time-consuming and complex. While the PerCa workflow (35) and its updated PCA-N variant (23) utilised an EvoSEP for in-line clean-up, we implemented STrap digestion with SPE in its absence.

Estimated protein input-to-output ratios revealed significant differences in post-digest recovery (Supplementary Table 1). Between neat methods, higher recovery was observed for STrap (83.1%, 29 µg) compared to SP3 (53.1%, 18 µg). For corona enrichment workflows, EnrichIST produced the highest post-digest yield (1.4%, 19 µg), with MagNet and ProteoNano showing comparable recoveries (0.20-0.32%; 5.6-11.4 µg). PerCA depletion resulted in lowest retrievals (0.12%, 4.17 µg), consistent with the PCA-N workflow (23), where a 5 µL input resulted in 1000 ng of material.

Peptides were analysed on two plasma-optimised nano-LC-MS methods on the Orbitrap Astral connected to a Vanquish Neo LC system (Figure 1A), titled ‘Discovery’ and ‘Throughput’. The ‘Throughput’ method used half the runtime (26 min vs. 50 min active gradient) and sample load (150 µg vs. 300 µg) compared to the Discovery setup.

### Workflow-specific enrichment and the trade-off between throughput and detection

Across LC-MS runs and methods, we identified a non-redundant total of 2,726 human proteins from 27,465 stripped peptides sequences, and 3,767 rat proteins from 40,215 stripped peptides. Precursor, peptide and protein counts for each workflow and LC-MS method are summarised in Supplementary Table 2.

Across the baseline (neat) methods, STrap yielded higher protein identifications in plasma, but slightly lower average peptide- and precursor-per-protein counts (Supplementary Table 2). SP3 exhibited the highest rate of missed cleavages across all workflows, including other on-bead digest methods with similar digestion times. In human biofluids, both neat workflows identified >1,000 proteins using the Discovery method. Using the Throughput method, STrap and SP3 identified 945 and 822 proteins, representing a 20% and 17% reduction in identifications, respectively (Supplementary Table 2; Figure 1C). In neat rat plasma, ∼1,500 proteins were identified with the Discovery method, with a similar 20% reduction observed when using the shorter Throughput method (Figure 1C). These increased identifications in rat plasma were due to a higher precursor count, with STrap yielding 3,771 and 2,041 more precursors in rat compared to human plasma for Discovery and Throughput, respectively.

MagNet-enriched biofluids performed similarly between bead chemistries and achieved the highest detection rates, with >20,000 precursors and ∼1,700 proteins identified in human biofluids, and >35,000 and ∼3,000 proteins in rat plasma using the Discovery method (Supplementary Table 2). With the exception of PerCa, all workflows yielded more identifications in rat plasma than in human biofluids. The greatest increase in human biofluids was 52% compared to 101.9% in rat (STrap vs MagNet HILIC; Discovery Method). Relative to neat plasma, all corona-enrichment methods increased protein identifications, with progressive gains observed from ProteoNano (19.8% in human, 35.3% in rat), to EnrichIST (34.5% in human, 67.6% in rat), and MagNet (42.9–51.9% in human, 89.0–101.8% in rat), the latter maintaining >90% reproducibility in rat plasma (Figure 1B). The reduced depth between Discovery and Throughput methods was also more pronounced in human biofluids for enrichment methods than to the rat, with 28.9% ± 2.6% reductions compared to 25.6% ± 2.6% in human and rat, respectively (Figure 1C).

With only a few exceptions, enrichment and depletion methods outperformed neat identifications. ProteoNano proved unsuitable for human serum, with a large bias towards plasma-unique identifications (Figure 1D; Supplementary Table 2). PerCa led to significant improvement in identifications in human biofluids, but had comparable depth to neat workflows in rat plasma. Intriguingly, PerCa had lowest peptideslzlperlzlprotein and precursorslzlperlzlprotein ratios across all workflows, with below-baseline precursors counts (Table S2). More specifically, although PerCa identified 45.2% more proteins than SP3 in human plasma, it detected 9.1% fewer precursors (and 33.9% less in equivalent rat comparisons). Similar precursor-to-protein ratios for PerCa occurred in Beimers *et al* (25) (Supplementary Figure 1). However, no workflow significantly exceeded the ratios observed in neat baselines, nor were these ratios substantially improved by longer LC-MS methods.

### Exploring biofluid, species and method-specific performance

Approximately one quarter of reproducibly identified proteins were shared between workflows (Figure 2). Notably, corona enrichment methods contributed significantly to unique protein identifications, with MagNet SAX and HILIC beads yielding 10% and 7.5% unique proteins in rat and human plasma, respectively (Discovery). In rat plasma, the top intersections of corona enrichment workflows encompassed 38% proteins not detected in neat baseline (Strap, SP3) or depleted (PerCa) workflows. In contrast, in human biofluids, the highest unique intersections were proteins shared between PerCa serum and plasma (127 proteins) and then uniquely detected in PerCa plasma (54 proteins), underscoring method-specific divergences between enriched and depleted proteomes. Although previous human benchmarking studies reported minimal differences between MS-enrichment workflows (25, 28), deeper coverage in rodent plasma highlighted pronounced method-specific distinctions.

**Figure 2.**
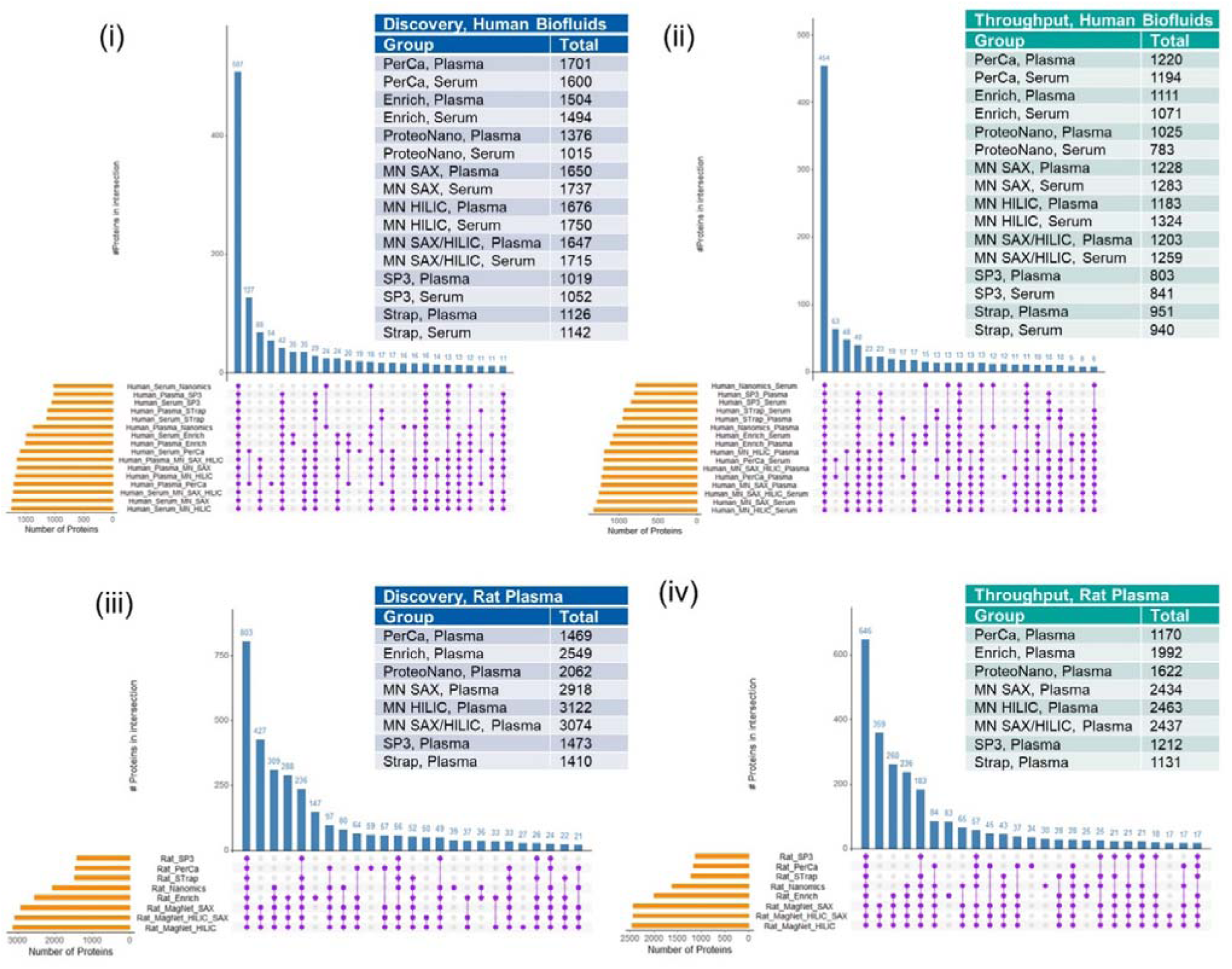
Evaluation of species- and method-specific identification and overlap. upSET intersection plots of reproducibly identified proteins (≥2 replicates) by workflow for: (i) human biofluids— Discovery method; (ii) human biofluids—Throughput method; (iii) rat plasma—Discovery method; (iv) rat plasma—Throughput method. An inset table reports the non-redundant totals of reproducible proteins per method and biofluid.

Principal-component analyses (PCA) (Figure 3A-B) confirmed clear segregation of workflows within each species, with both PC1 and PC2 variation driven predominantly by workflow. Intriguingly, the PerCa workflow clustered with human baseline workflows along PC1, while ProteoNano was closest to the rat baseline workflows along PC1, with PerCa separated on the PC2. By contrast, the three MagNet variants (HILIC, SAX and their combined HILIC/SAX protocol) occupy almost identical coordinates in both species, separated from all workflows along PC1, consistent with this axis capturing differences between EV-specific and secreted plasma proteomes (34).

**Figure 3.**
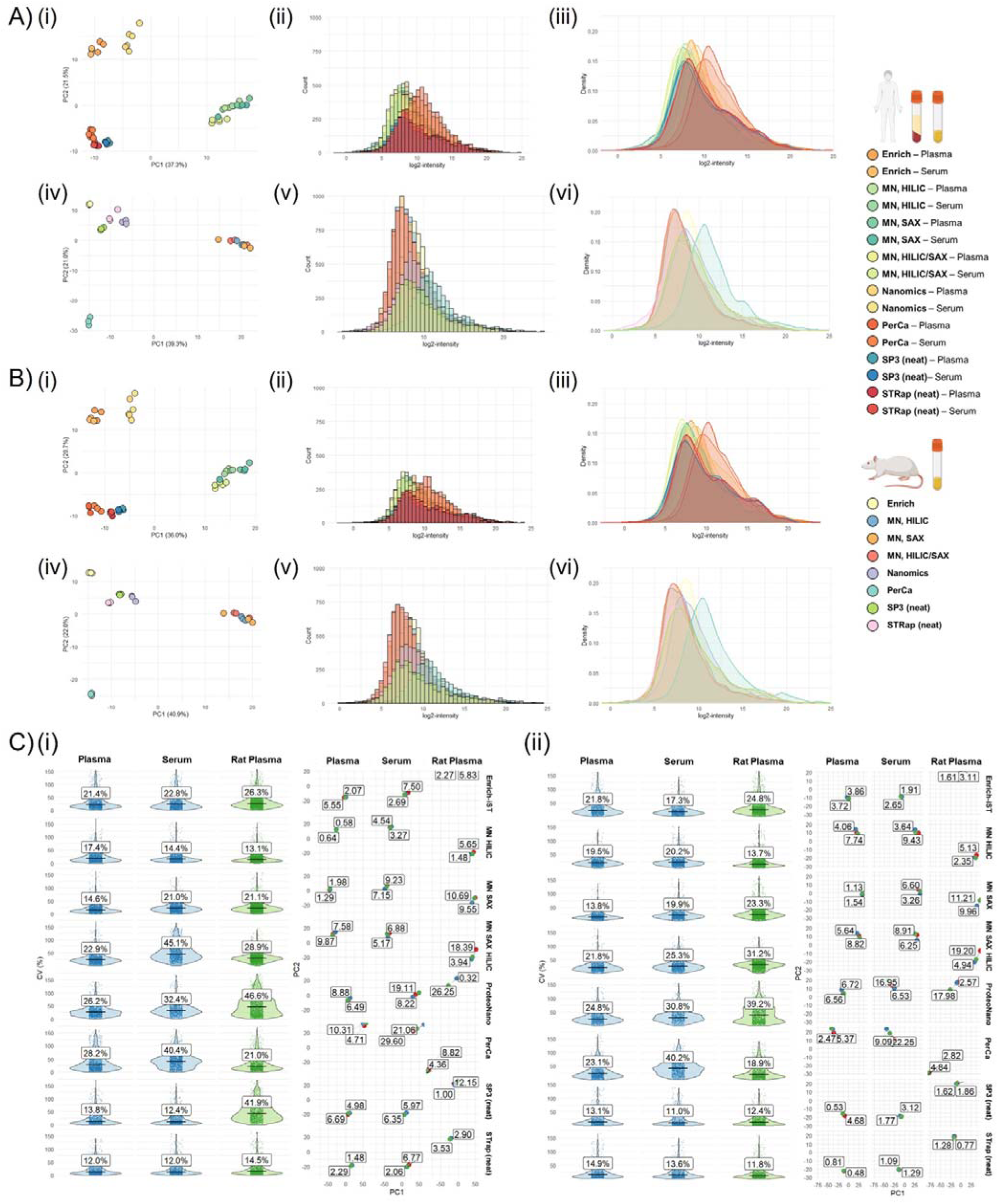
**Evaluation of species- and method-specific performance metrics.** (A) Charts for the **Discovery method** in human biofluids (i – iii) and rat plasma (iv – vi) and (B) Charts for the **Throughput method** in human biofluids (i – iii) and rat plasma (iv – vi). Includes (left through right) PCA of log2-transformed intensities, binned histograms (protein counts per intensity bin), kernel density curves. (C) Left: violin plots of univariate variance (CV%) of **untransformed** intensities by workflow and biofluid, computed on reproducible proteins (≥2 replicates); median CV% annotated on each violin. Right: multivariate variance via a single global PCA fit on centred and scaled protein profiles across all samples; panels show PC1 vs PC2 with shared axes for cross-method comparison. Points denote replicate injections (1–3), coloured by replicate; within each method, arrows trace 1→2 and 2→3, with labels indicating Euclidean distance between adjacent replicates in the PC1–PC2 plane. Data is shown for the (i) Discovery method and (ii) Throughput method.

The notoriously challenging dynamic range of plasma was evident in neat-plasma density plots, which showed a pronounced right tail in the protein intensity distribution. Consequently, successful compression of dynamic range was expected as the reduction in kurtosis, and the leftward shift for protein counts in lower intensity ranges (Figure 3A-B). The higher albumin concentration in humans (39), along with other abundant suppressive proteins (40), was evident in a more pronounced kurtosis in neat human compared to rat plasma. The latter had reduced suppression of lower abundance proteins, and improved detection across all workflows, with doubling of protein counts in lower intensity ranges for enrichment workflows, notably for MagNet. While human samples also showed reduced right-tail skewing (Supplementary Figure 2), improvements in low-intensity protein counts were less pronounced. Interestingly, PerCa depletion reduced kurtosis in both species but caused a rightward shift in the whole intensity distribution.

Given the deeper coverage of low-abundance proteins in rat plasma, we examined trends in the coefficients of variation (CV) across the various workflow and species (Figure 3C). We found no evidence that proteome depth or species affected quantitative reproducibility. To assess technical variation, we examined both univariate CV% and multivariate PCA distances. This allowed us to distinguish between handling issues – where a single outlier replicate skewed PCA clustering and increased CV – from random noise (high CV%, low PCA distance) and structured noise (moderate CV%, high PCA distance). Most elevated CVs in workflows such as PerCa and ProteoNano were due to a single dissimilar replicate, which could be mitigated through optimised, automated preparation (21, 23, 28). General trends included a modest reduction in CV with the Throughput method, likely due to under-representation of low-abundance proteins with shorter analysis times, and a more pronounced decrease in neat preparations, reflecting the inherent stability of high-abundance plasma proteins and fewer handling steps. However, low and consistent levels of variation across the 9 replicates and 3 inputs was observed in MagNet HILIC and EnrichIST, crediting these as most confidently biofluid- and species-agnostic enrichment methods.

### Workflow overlap with plasma reference and functional databases

Highly abundant plasma proteins are known to mask and suppress detection of low-abundance targets (11). As such, we assessed changes in log_2_ intensity of the top 100 abundant plasma proteins, using STrap neat plasma as the baseline (Discovery data; Figure 4A). Notably and as expected, the deviation between SP3 and STrap intensities for the top 100 was limited, given both represent neat plasma proteomes.

**Figure 4.**
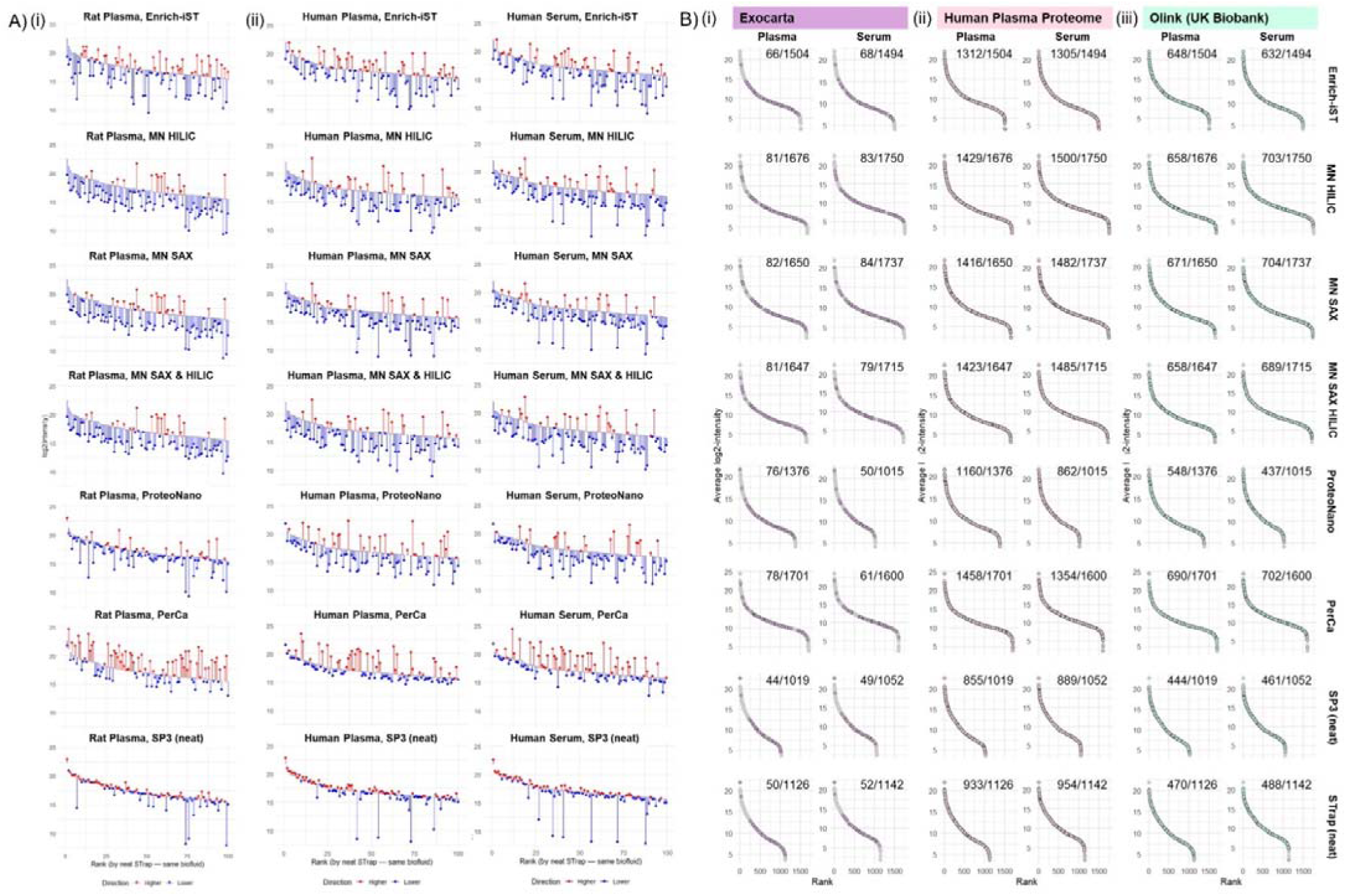
**Assessment of workflow enrichment (Discovery method).** (A) Rank-abundance plots of the top 100 plasma proteins, using STrap as the baseline, for (i) rat plasma, (ii) human plasma, and (iii) human serum. Baselines are defined by the average log2 replicate intensities from STrap within each biofluid. For each protein, the average log2 intensity in each workflow is plotted as a point; a segment connects the STrap baseline to the workflow value, coloured red if higher than baseline and blue if lower. (B) Rank-abundance plots by workflow and human biofluid showing average log2 intensities for reproducibly identified proteins (≥2 of 3 replicates). Proteins present in reference sets are highlighted, with total detections labelled in the top-right of each panel. Reference sets: (i) ExoCarta Top 100 (purple), (ii) HPP 2021 build (10) (4394 proteins; pink) and, (iii) Olink 3K targets (41) (2941 proteins; teal).

Corona enrichment workflows, particularly MagNet, successfully reduced the intensities of nearly all top 100 plasma proteins, with widespread depletion also observed in EnrichIST (Figure 4A). Intriguingly, while widespread depletion was also observed in ProteoNano, the effect was less pronounced in rat plasma, indicating species-specific differences. Unexpectedly, PerCa depletion resulted in increased intensities for abundant plasma proteins, likely reflecting a global shift in log_2_ intensity distributions (Figure 3A-B). Change in intensities across the entire baseline are shown in Supplementary Figure 3, with counts of proteins above or below baseline levels detailed in Supplementary Table 3.

While these findings demonstrate that workflows unequivocally reshaped the native plasma proteome (Figure 4A), we further evaluated whether these changes were due to compositional shifts (i.e. through enrichment of circulating-EVs (34)), or simply improved detection through dynamic range compression. To explore this, we compared proteins identified in human biofluids (Discovery data) against established references and databases (Figure 4B), including the top 100 EV markers from ExoCarta, proteins from the Human Plasma Proteome (HPP) (10), and plasma markers characterised from the UK biobank using Olink (41).

As expected, EV-specific markers were most frequently detected in MagNet workflows, and least in the two neat baseline methods (SP3 and STrap). EV marker abundances also increased in rank within MagNet datasets, concordant with EV enrichment. After neat workflows, EnrichIST showed the lowest detection of EV markers, even lower than ProteoNano and PerCa, which credited the corona was not specific for membrane bound vesicles. Intriguingly, EV marker numbers were markedly lower in PerCa-depleted human serum (n = 61) than in plasma (n = 78), despite the latter containing only 101 additional reproducibly detected proteins. Otherwise, numbers of EV markers were similar between the two human biofluids.

For HPP and Olink proteins, detection scaled with proteome depth achieved by each workflow. HPP coverage in the datasets was high, ranging from 82.7% (STrap Plasma, n = 933) to 87.4% (EnrichIST Serum, n = 1,312), as expected given both are derived from MS data. Conversely, the number of MS-detectable Olink-markers was far lower ranging from 39.8% to 43.9%, with the highest counts just over 700 proteins in serum samples processed with MagNet HILIC, MagNet SAX, and PerCa. Several benchmarking studies (7, 42) have also reported limited overlap between MS and Olink proteomics, consistent with their differing principles – antibody affinity versus native protein abundance – resulting in complementary dynamic ranges and detection.

### Exploring cell- and tissue-specific origins of depleted and enriched proteomes

To investigate compositional changes, we mapped our identified proteins in enriched and depleted proteomes to a recent large-scale, plasma-specific tissue and cell atlas comprising 18 tissues and 8 immune cell types, covering 18,430 proteins across 688 maps (5) (Figure 5A). From our dataset, we matched 1,578 human proteins to 234 maps (Supplementary Figure 4). To simplify interpretation, we focussed on tissue-specific, cell-specific and common plasma maps, which retained 1,019 proteins from 29 maps. Similar to HPP, the number of mapped proteins scaled relative to proteome depth, with highest marker counts observed in MagNet-enriched serum (n = 740 in HILIC, n = 710 in SAX), although marker increase in serum were not generalisable.

**Figure 5.**
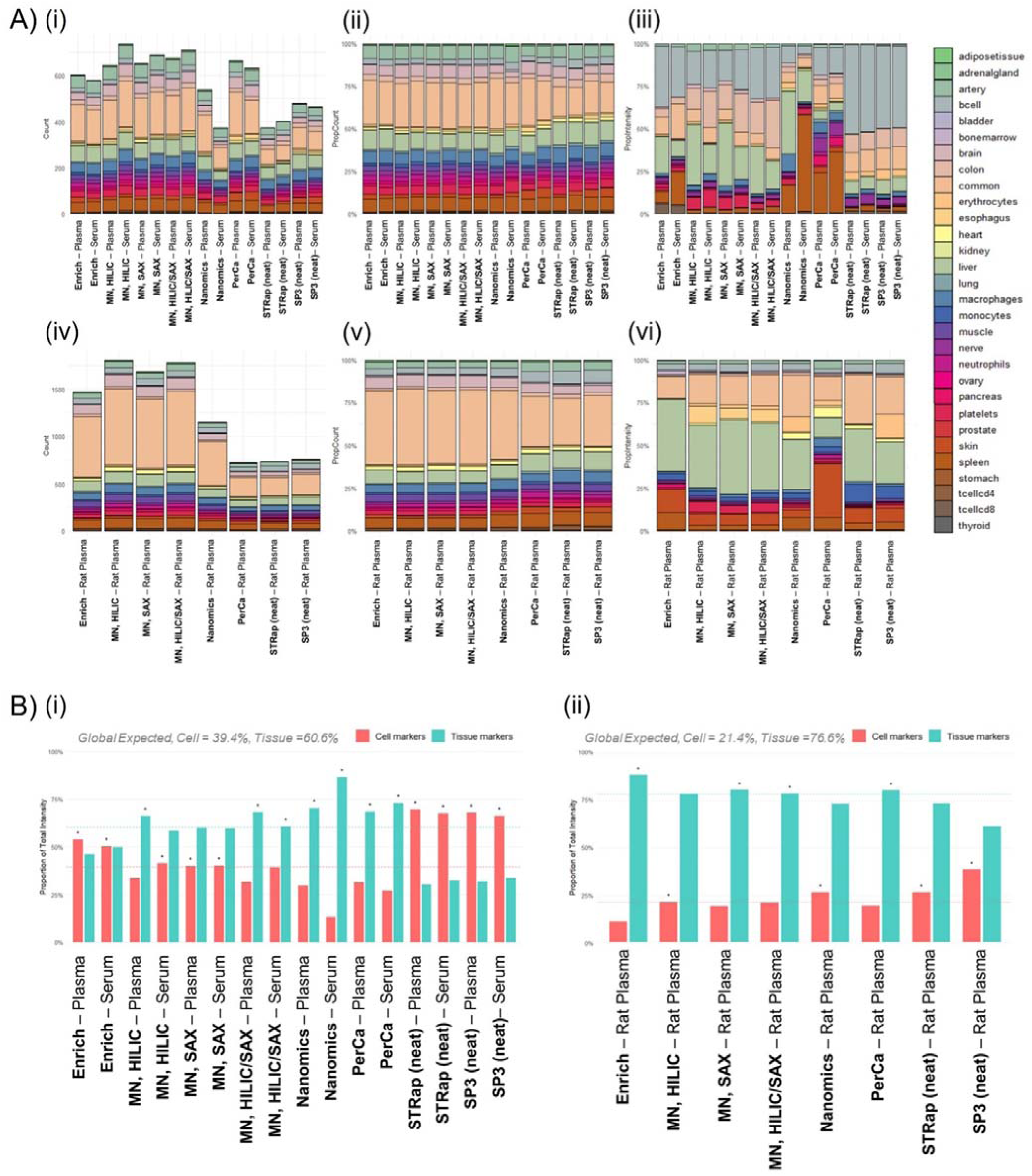
**Cell and tissue distribution of plasma proteins by workflow (Discovery method).** (A) Human biofluids: stacked bar charts summarising, for each workflow, (i) raw protein counts per global label, (ii) percent of protein counts per label, and (iii) percent of total intensity per label. Counts are restricted to unique proteins per method x label with a single, unambiguous cell or tissue assignment in the atlas (5). Intensities are summed within method × label after back-transformation from log2 and expressed as a proportion of each method’s total intensity (0–100% scale). (B) Rat plasma (human-mapped orthologs): summaries as in (A), computed after mapping rat proteins to their human orthologs. (C) Cell-versus tissue-marker enrichment by workflow for (i) human biofluids and (ii) rat plasma. Grouped bars show the proportion of total marker intensity assigned to cell markers and to tissue markers for each method. Dashed horizontal lines indicate expected background proportions estimated from pooled intensities across all samples. Enrichment is assessed with one-sided exact binomial tests (rounded summed intensities treated as counts), with Benjamini–Hochberg adjustment across methods; asterisks denote adjusted p < 0.05 for either cell- or tissue-marker enrichment.

Among neat baselines, SP3 detected fewer specific markers than STrap (Figure 5A; Supplementary Figure 4). As expected, MagNet-enriched biofluids yielded the greatest enrichment of immune cell markers, notably from macrophages, monocytes, and T cells, confirming circulating EVs as the preferred workflow for profiling immune-cell communication. Detection of distinct markers within the same map was observed. For example, 71 brain-specific proteins were identified across all workflows, ranging from 23 (STrap, plasma) to 51 (HILIC, serum). Of these, 68 were detected between HILIC-enriched and PerCa-depleted workflows, but only 38 (55.9%) were shared. Furthermore, PerCa had unique tissue-specific coverages, including 21/23 esophageal markers, 5/5 of adrenal markers and 40/40 of skin markers.

When comparing tissue- and cell-specific markers intensities, we observed dramatic compositional shifts (Figure 5A). As expected, summed intensities of B-cell markers (n = 23), dominated by immunoglobulins, accounted for half of the total neat biofluid intensities, but were reduced by more than half in other workflows. Platelet-specific markers were most abundant in MagNet workflows, reinforcing recent concerns about potential bias for pre-analytical contamination (27). However, only ProteoNano samples tested positive for platelet contamination using *Gao et al* (17) NP-specific Biaze tool (Supplementary Figure 5). MagNet workflows also showed high proportions of colon and arterial markers, with HILIC yielding the highest proportion of brain-specific proteins. PerCa again revealed high tissue-specific proportions, including spleen-specific proteins (reflected to a lesser extent in ProteoNano and then EnrichIST), in addition to skin, pancreas, nerve, muscle and lung markers.

To determine whether specific cell- or tissue-specific markers were disproportionately represented in each workflow, we performed one-sided binomial tests comparing the observed intensity fraction of each maker against its global background. Cell-enriching workflows included neat workflows, EnrichIST and MagNet SAX, while tissue-enriching included PerCa and ProteoNano and MagNet SAX-HILIC combination. MagNet HILIC was the only workflow with differential enrichment between plasma and serum for tissue- and cell-enriching, respectively.

Given these distinct enrichment profiles, we extended our analysis to rat proteomes using homology mapping. We matched 2,077 of 3,663 rat orthologs to human identifiers, covering 30 cell- and tissue- specific maps. Intriguingly, only 491 proteins (21.1%) were shared between the mapped rat orthologs (1,801) and human markers (1,019). As in the human datasets, marker counts scaled with proteome depth, and were observed to be highest in EV-specific workflows, with 1,808 markers detected in MagNet HILIC (Figure 5B). PerCa depletion on rat plasma showed strong enrichment for esophageal, lung, and skin markers, and distinct markers within the same maps were observed. As noted previously, 148 of 152 rat-mapped brain-specific markers were detected across EnrichIST (96), MagNet-HILIC (120) and PerCa (42) workflows, with only 24 (16.2%) shared and 18, 39 and 5 were unique to each workflow, respectively.

Differing inter-species dynamic ranges were reflected in decreased compositional proportions of B-cell proteins in rat plasma to human (Figure 5A). As in human biofluids, MagNet-enriched rat samples showed elevated platelet marker intensities, while PerCa retained high proportions of spleen and skin markers, and also enriched for heart and arterial proteins. When applying binomial tests, PerCa remained tissue-enriching, while neat workflows were cell-enriching. Notably, all corona-based workflows – except MagNet SAX-HILIC – reversed their enrichment category between human and rat datasets (Figure 5B). We suggest that these shifts likely reflect interspecies differences in dynamic range and the increased detection of lower-abundance proteins.

### Statistical evaluation of differentially abundant proteins between workflows

We used the DIA Analyst platform for statistical evaluation of differentially abundant proteins (DAPs). Nearly all proteins were observed to be significantly regulated in at least one pairwise comparison within rat plasma (Figure 5, Discovery 97.7%, Throughput 96.6%), and human biofluids (Figure 6, Discovery 99.7%, Throughput 99.6%). Clustered heatmaps of proteins revealed distinct blockwise-patterns of workflow-specific enrichment in both species (Figure 6A and 7A), consistent with previous MS-based plasma benchmarking studies (25, 26, 28), particularly between EV-specific and secreted proteomes.

**Figure 6.**
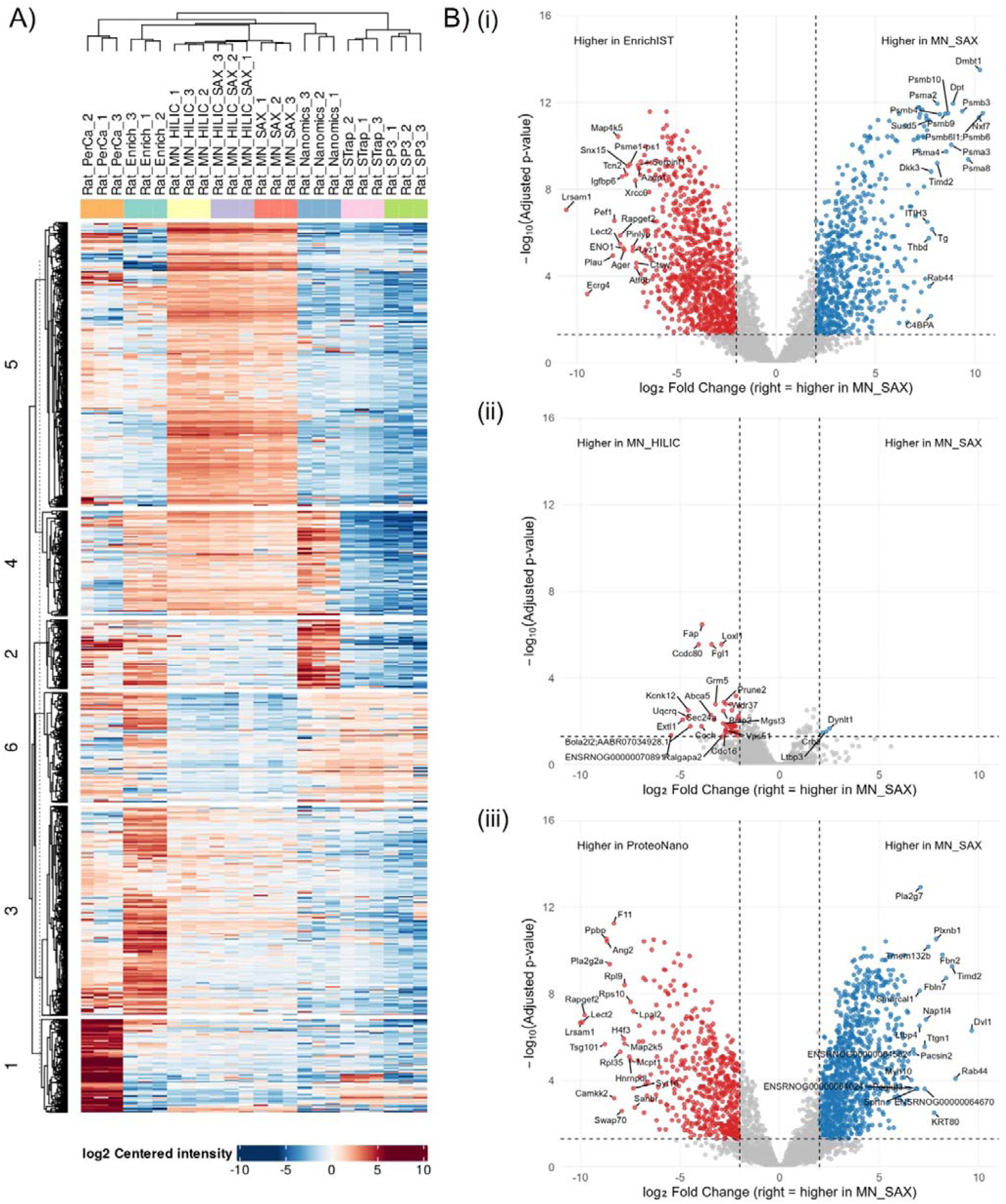
**Differentially abundant proteins in rat plasma across workflows (Discovery method).** (A) Hierarchical clustering heatmap of proteins (rows) across workflow x replicate samples (columns) generated in DIA Analyst. Values are log2 mean-centred intensities per protein across all samples; blue indicates below-mean abundances and red above-mean abundance (scale bar, −10 to +10). Columns and rows are hierarchically clustered (dendrograms shown); numbered brackets at left denote six major protein clusters obtained by cutting the row dendrogram. (B) Volcano plots for pairwise comparisons among corona-based enrichment workflows: (i) MagNet-SAX versus EnrichIST, (ii) MagNet-SAX versus MagNet-HILIC, and (iii) MagNet-SAX versus ProteoNano. Axes show log2 fold change (x) versus −log10 (adjusted p) (y); positive x (right side) indicates higher abundance in MagNet-SAX. Dashed vertical lines indicate fold-change thresholds (log2FC = 2) and the dashed horizontal line indicates the significance threshold (FDR-adjusted p = 0.05). Points are coloured by significance category: higher in MagNet-SAX (blue), higher in the comparator (red), and not significant (grey). The 20 most extreme significant proteins on each side (largest log2FC, then lowest p) are labelled by gene name

**Figure 7.**
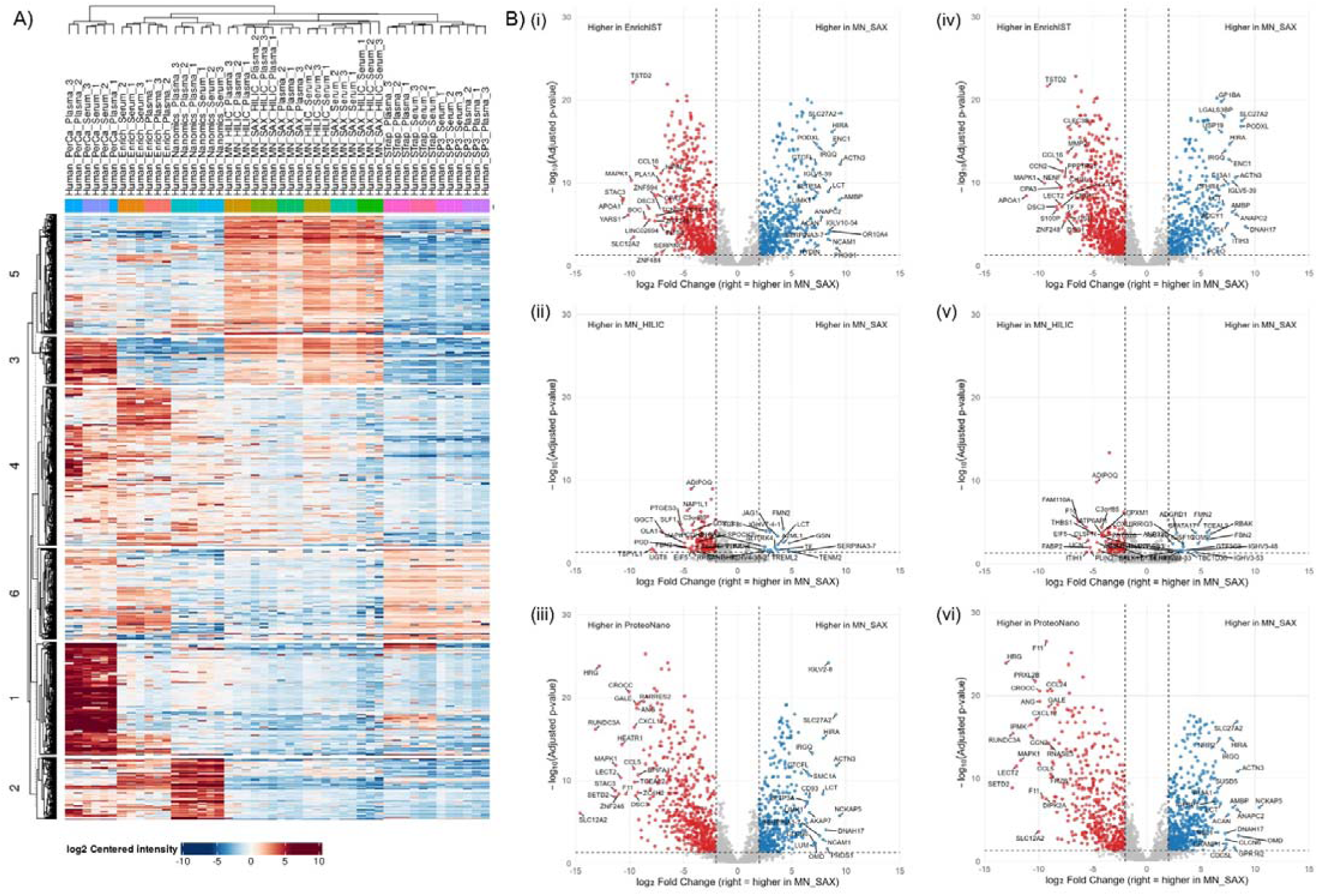
**Differentially abundant proteins in human plasma and serum across workflows (Discovery method).** (A) Hierarchical clustering heatmap of proteins (rows) across workflow x replicate samples (columns) generated in DIA Analyst. Values are log2 mean-centred intensities per protein across all samples; blue indicates below-mean abundances and red above-mean abundance (scale bar, −10 to +10). Columns and rows are hierarchically clustered (dendrograms shown); numbered brackets at left denote six major protein clusters obtained by cutting the row dendrogram. (B) Volcano plots for pairwise comparisons among corona-based enrichment workflows: (i) MagNet-SAX versus EnrichIST (Plasma), (ii) MagNet-SAX versus MagNet-HILIC (Plasma), (iii) MagNet-SAX versus ProteoNano (Plasma), (iv) MagNet-SAX versus EnrichIST (Serum), (v) MagNet-SAX versus MagNet-HILIC (Serum), and (vi) MagNet-SAX versus ProteoNano (Serum). Axes show log2 fold change (x) versus −log10(adjusted p) (y); positive x (right side) indicates higher abundance in MagNet-SAX. Dashed vertical lines indicate fold-change thresholds (log2FC = 2) and the dashed horizontal line indicates the significance threshold (FDR-adjusted p = 0.05). Points are coloured by significance category: higher in MagNet-SAX (blue), higher in the comparator (red), and not significant (grey). The 20 most extreme significant proteins on each side (largest log2FC, then lowest p) are labelled by gene name.

When comparing human serum to plasma (Supplementary Figure 6-7), fewer DAPs were observed between these biofluids than between human workflows (Supplementary Figure 9-10). Intersections of workflow DAPs in serum versus plasma identified up to 304 DAPs in PerCa (the lowest was 68 DAPs in SP3), as expected based on numerous PerCa-unique proteins at the identification-level (Figure 2A). Only six DAPs between plasma and serum were shared across workflows, four of which were fibrinogens (FGA, FGB, FGB.1, FGG.1) higher in plasma. Outside the shared set, most serum-plasma DAPs were workflow-specific (Supplementary Figure 7), suggesting that QC markers for evaluation of preparation effects (e.g., collection tubes) and contaminants (e.g., coagulation) may require either workflow-specific marker panels or marker-level reference ranges.

In rat plasma, MagNet workflows and EnrichIST resulted in the greatest number of DAPs compared to neat baselines (Supplementary Figure 8). The largest intersections of DAPs were observed between the three MagNet bead chemistries, followed by unique DAPs in EnrichIST in both the Discovery and Throughput method (Supplementary Figure 11-12). Slightly lower numbers of DAPs were observed for PerCa and ProteoNano, and only a few between neat baselines. DAP distributions between neat baselines were consistently biased towards STrap, reflecting SP3’s lower intensities in rank-abundances (Figure 4A).

Hierarchical clustering of rat DAPs revealed discrete workflow-associated enrichment patterns (Figure 6A). The largest cluster encompassed MagNet EV-enriched proteomes, with protein-protein interaction (PPI) networks that highlighted a dominant proteostasis complex, including the proteosome, lysosome and chaperones, alongside extracellular matrix and membrane-trafficking/endosomal terms (Supplementary Figure 13). This is consistent with charge-driven capture mechanisms that selectively recover ECM fragments and EVs (34). The PerCa cluster was enriched for skin-associated terms and extracellular filament/cytoskeletal components, mirroring the signatures observed within the cell/tissue atlas (Figure 5B). EnrichIST-enriched proteins reinforced the differences in corona-chemistries differences based on cellular amino-acid and small-molecule metabolism (Supplementary Figure 13), with a similar MagNet signature of folding, endosomal transport, and structural terms (ECM, collagen, actin signature).

A shared corona-enrichment workflow cluster further reinforced overlaps MagNet, EnrichIST and ProteoNano. This cluster included PPI networks involved in RNA-rich particles and ribonucleotide protein (RNP) complexes (Spliceosome) (Supplementary Figure 14), with GO enrichment confirming associations with EV proteins (including platelets), clathrin-mediated endocytosis, vesicle acidification, and proteostasis. Together, these findings suggest that rat corona workflows consistently enriched a shared subset of vesicular-derived proteins, where dynamic range compression post-enrichment enhanced detection of low-abundance secreted proteins and EV components (Figure 4B).

Human biofluids exhibited less workflow-specific clustering (Figure 7A), with only MagNet and PerCa forming enrichment-specific clusters. As the clustering included in both serum and plasma, matrix-driven signals (notably clotting and platelet-derived EVs) likely masked and suppressed workflow-specific clustering. The human MagNet cluster highlighted ECM organisation and cell adhesion, glycosaminoglycan metabolism, haemostasis and wound healing (Supplementary Figure 15), diverging from the proteostasis-dominated rat MagNet cluster (Supplementary Figure 13-14). In contrast, enriched human PerCa proteins mirrored the signature observed in rat plasma, with skin and epidermal functional terms, including intermediate filament organisation, keratinocyte and epidermal differentiation, and unique terms for peptide cross-linking and endopeptidase regulation.

Lastly, given that corona-enrichment comprised half the tested workflows, we examined the DAPs in paired tests between the four major corona methods (MagNet SAX, MagNet HILIC, EnrichIST and ProteoNano). In the rat plasma (Figure 6B), DAPs between MagNet SAX and HILIC were minimal (49 DAPs, 0.001%), but intriguingly higher in human biofluids (150 plasma DAPS, 5.7%; 110 serum DAPS, 4.2%) (Figure 7B), with MagNet HILIC consistently yielding more DAPs. Low DAP numbers were reproducible across inter-MagNet comparisons in rat (< 2%), while higher DAP numbers were observed in human comparisons, particularly in serum (Figure 7B). However, distinct compositional differences were evident between MagNet, EnrichIST and ProteoNano coronas. In rat plasma, DAPs between EnrichIST and MagNet SAX were higher (1576, 43.5%) than ProteoNano (1516, 39.3%), although the highest inter-corona DAPs were similar for MagNet HILIC with EnrichIST (1721, 47.5%) and ProteoNano (1709, 47.2%). In human biofluids, more than half of all proteins were differentially abundant between EnrichIST and MagNet SAX/HILIC (1453, 55.4%; serum) and ProteoNano and MagNet SAX/HILIC (1418, 54.0%; serum). Collectively, these data reaffirm pronounced, corona-chemistry-dependent shifts in the enriched plasma proteome, where the species context magnified these effects.

## Discussion

Liquid biopsies offer a means to capture biomarker molecules or signatures in circulation. As demonstrated here, recent advances in proteomic workflows have substantially reduced MS-related challenges in measuring plasma protein markers, and these workflows are now applicable across both clinical cohorts and preclinical models. In the case of neat plasma, where dynamic range suppression is most severe, our findings underscore the remarkable progress of next-generation mass analysers, readily surpassing the 1,000 protein mark when combined with plasma-optimised LC methods (Figure 2). We also observed depth was improved by adjusting the MS method (Figure 1C), whereby a longer LC-MS analysis time particularly enhanced enrichment workflows, as shown both in this study and by others (28). However, despite evaluating eight sample-processing workflows, three biofluids across two species, and two plasma-optimised MS methods, this study still interrogates only a fraction of the available parameter space. Nevertheless, it reinforces the emerging consensus that clinical proteomic workflows should be selected using context-aware, data-driven criteria.

Our protein identifications in human neat plasma exceeded those reported in recent benchmarking studies utilising the Astral mass spectrometer (25, 28), even when filtered for reproducibility (Figure 1B). Intriguingly, our two neat plasma workflows did not perform equally (Figure 3, Figure 4), suggesting underlying factors that may explain the variability in plasma results across studies (25, 26, 28). In contrast, some of our enriched-proteomes identifications were also lower than expected (28). This was somewhat expected, given our use of pooled samples across multiple healthy donors, which tends to average out low-abundance, donor-specific features. Healthy plasma furthermore lacks pathology-driven signals such as infection- or inflammation-related signatures, and previous studies have indeed shown that technical replicates of healthy samples show reduced identifications compared to clinical cohorts (25). Furthermore, specific workflows may perform better in certain contexts. For example, under pro-inflammatory conditions (e.g., viral sepsis or COVID-19), EV-enrichment may confer proportionally greater improvements due to disease-driven changes in EV cargo. By the same token, incorporating more low-abundance proteins and individual variability will also elevate CVs, such that reported values here must also be considered in relative terms, and users must be mindful that quantitative consistency remains as important as depth (26).

Comparing across the expanding benchmarking scholarship is increasingly challenging. Studies differ in their use of plasma sources, workflows, and plasma collection protocols (27), which credits recent calls for standardisation through reference samples or spike-ins (43). However, as these discussions evolve, there is an opportunity to ensure these that these methods for standardisation will support cross-species evaluation. In non-human species, MS remains the best technology for pre-clinical biomarker discovery, noting that aptamer- and antibody-based technologies such as Olink or SomaScan (44)) offer limited coverage for mice only. Our results highlight that clinical proteomics innovations can be successfully applied to preclinical models, where performance of truly species-agnostic workflows was only influenced by the species-specific dynamic range. Moreover, in small-mammalian rodent models, where high-abundance proteins are less suppressive than in humans, MagNet and EnrichIST performed particularly well, with EnrichIST on the Astral eclipsing numbers reported by its manufacturer (45). While ProteoNano also performed well in rat plasma, caveats emerge given its prescribed use for human plasma only, the fact we observed reduced depletion efficacy of the top 100 proteins in rat plasma (Figure 4A), and clear unsuitability for human serum (Figure 1D). However, re-tuning ProteoNano for other species is likely more feasible than re-engineering aptamer platforms, underscoring the need to include non-human biofluids in the plasma-technology discourse given they are a crucial proving ground for human trials.

Our results also contribute to ongoing discussions surrounding prelzlanalytical biases in deep proteomics and bead-based workflows (17, 27), particularly from PMBCs and platelets. Recent studies found that EV-enrichment (MagNet-SAX) was highly sensitive to technical and cellular contamination (27), whereas depletion via PCA-N was largely unaffected. Consistent with this, our data showed platelet-specific marker counts and intensity proportions in PerCA resembled neat workflows and were highest in MagNet (Figure 4). This is unsurprising as EV-associated proteins represent only a small fraction of the plasma proteome, and perchloric-acid precipitation is a harsh denaturing step that would not preserve intact EVs, thereby redistributing vesicular cargo into the bulk protein pool. However, we propose that the extent and nature of this co-enrichment reflect both ligand-corona specificity (biased capture toward subproteomes enriched for technical artefacts) and the workflow’s enrichment factor, which increases both genuine and artefactual signals. In this vein, we note EnrichIST only exhibited a modest increase in platelet signal (Figure 5) and fewer EV-specific markers than PerCA, and much lower than MagNet (Figure 4A). Furthermore, among the five corona workflows, platelet contamination was only observed in one of our NP systems (Supplementary Figure 5). This finding appears more in line with Guo *et al* (17), where the degree of blood-specific contaminants varied between NP chemistries.

Contrary to the notion that identification rates in bead-enrichment workflows indicate contamination rather than dynamic range improvement, our results in rat plasma revealed many more tissue-specific markers in deeper, bead-enriched proteomes compared to humans, including heart, brain and pancreatic markers. Of course, we do not advocate for complacency. Artefacts will dramatically increase in improperly processed plasma such as incomplete clotting or processing at high *g*-force. In these cases, contamination may be assessed by indexation of a workflow’s specificity for cell-specific markers (the source of significant pre-analytic biases (3, 27)), or mitigated by analysing only tissue-specific markers if cohort pathology permits, given corona-enrichment methods were not exclusively cell-enriching (Figure 5B), nor were EV markers equally enriched (Figure 4B).

These differences underscore another important observation: proteomic depth was not uniform across workflows, it was redistributed. Our analyses of database-specific (Figure 4) and marker-specific (Figure 5) proteins revealed that depletion approaches enhanced coverage of mid-to-low abundance proteins where many tissue-specific and leakage proteins are readily detectable. In contrast, ligand-mediated enrichments favoured discrete, corona-defined subsets. In the case of the latter, even coronas from the same manufacturer with related chemistries (HILIC vs SAX) yielded hundreds of significantly regulated proteins, while comparisons across different coronas revealed over 1000 (Figure 7). The dynamic range effect also differed by species, where reduced suppression in rat plasma broadened accessible depth, specific maker detections, and amplified workflow-specific divergences (Figure 6). Accordingly, method selection should hinge on a workflow’s ability to enrich the proteins that matter, either through suitable dynamic-range access or targeted enrichment of key relevant subproteomes.

Plasma remains a dynamic and inherently complex biofluid for proteomic assessment, not just because of its high-abundance proteins, but because of its compositional plasticity across physiology and pathology. In these contexts, plasma proteomes diverge profoundly from healthy baselines and from each other. As clinicians and researchers demand rigorous clinical proteomics, species, symptomatology, and systemic responses within a given cohort must inform methodological choices: what is most likely to change, and which workflow is best poised to capture it? This isn’t necessarily an admission that methods are “biased” toward certain proteins, though we must accept that some workflows are sensitive to tube-related and other technical artefacts. Rather, it’s an acknowledgement that different workflows illuminate different plasma proteomic subsections and subsystems, the relevance for each can only be defined by that pre-/clinical cohort. There is no one-size-fits-all solution, only the opportunity (or the pitfall) to choose the best fit for the day, the destination, and the dress code.

## Acknowledgements

We acknowledge funding support from the Medical Research Future Fund (NCRI000108; R.B.S). All proteomic analyses were conducted at the High-Precision Biomarker Discovery node of the Monash Proteomics and Metabolomics Platform, where infrastructure enabled by Bioplatforms Australia (BPA) and the National Collaborative Research Infrastructure Strategy (NCRIS) was utilised. Additional support was provided by Monash eResearch capabilities, including Research Data Storage and Nectar Research Cloud.

## Data Availability

All raw files and their respective Spectronaut search outputs can be accessed via PRIDE with the identifier PXD072131.

## Author Contributions

SJEC: Conceptualization; Data curation; Formal analysis, Investigation, Methodology, Visualisation, Project Administration, Writing - original draft; and Writing - review & editing

JRS: Conceptualization, Methodology, Investigation, Data Curation, Writing - review & editing

IH: Conceptualization, Methodology, Investigation, Data Curation, Writing - review & editing

KKS: Conceptualization, Methodology, Investigation, Data Curation, Writing - review & editing

HL: Methodology, Writing - review & editing DHM: Methodology, Writing - review & editing

SB: Methodology, Investigation, Data Curation, Writing - review & editing ET: Resources, Methodology, Investigation, Writing - review & editing IA: Resources, Investigation, Writing - review & editing

TJOB: Resources, Investigation, Writing - review & editing PF: Resources, Methodology, Writing - review & editing

RBS: Conceptualization, Formal Analysis, Resources, Supervision, Project Administration, Funding Acquisition, Writing - original draft; and Writing - review & editing.

## Abbreviations

2-CAA: 2-chloroacetamide
ACN: Acetonitrile
DAP: Differentially Abundant Protein
DDA: Data Dependent Acquisition
DIA: Data Independent Acquisition
ECM: Extracellular Matrix
EDTA: Ethylenediaminetetraacetic Acid
EV: Extracellular Vesicle
FA: Formic Acid
GO: Gene Ontology
HILIC: Hydrophilic Interaction Liquid Chromatography
LC: Liquid Chromatography
MS: Mass Spectrometry
NP: Nanoparticle
PCA: Perchloric Acid
PPI: Protein-protein Interaction
SAX: Strong Anion Exchange
SPD: Samples Per Day
TCEP: Tris (2-Carboxyethyl) phosphine
TFA: Tri-fluoro acetic acid

