## Supplementary Data Files for "‘If the shoe fits?’ – technical nuances of plasma proteomic workflows in clinical and preclinical contexts"

#### Supplementary Information:

**Supplementary Table 1. Workflow logistics, inputs, and incubation/digestion steps.** For each workflow, the table lists input biofluid and plasma volume (µL), the estimated protein input (µg; assuming 70 µg per 1 µL plasma), key incubation steps (lysis, reduction, alkylation), digestion conditions/durations, and the minimum number of processing days. The total plasma volume used per workflow is reported. For the neat workflow, efficient lysis and complete reduction/alkylation require 2 µL of plasma; however, only 25% of the lysate is taken forward to digestion **(**35 µg). SPE was performed on Day 2 for all workflows, aligned with completion of digestions (including those requiring overnight incubation). Processing-day counts reflect optimal scheduling if this was not required.

| Method | Input (µL) | Biofluid Incubation | Incubation | Digest | Processing Days *Not including SPE* | Input (µg) *1 µL plasma = 70 ug* | Output (µg) *Avg from Nanodrop* | % recovery **input/output* |
|---|---|---|---|---|---|---|---|---|
| Strap (neat) | 2 | NA | NA | Overnight (Trypsin/LysC) | 2 | 35 * | 29.08 | 83.08 |
| SP3 (neat) | 2 | NA | NA | On bead, 1-2 hrs (Trypsin/LysC) | 1 | 35 * | 18.60 | 53.13 |
| MagNet –SAX | 50 | Dilute (buffer), on bead incubation (30 min), washes x 3 (15 min) | 60 min | On bead, 1-2 hrs (Trypsin/LysC) | 1 | 3500 | 11.36 | 0.32 |
| MagNet –HILIC | 50 | Dilute (buffer), on bead incubation (30 min), washes x 3 (15 min) | 60 min | On bead, 1-2 hrs (Trypsin/LysC) | 1 | 3500 | 7.25 | 0.21 |
| MagNet –SAX + HILIC | 50 | Dilute (buffer), on bead incubation (30 min), washes x 3 (15 min) | 60 min | On bead, 1-2 hrs (Trypsin/LysC) | 1 | 3500 | 7.69 | 0.22 |
| Enrich | 20 | Neat Biofluid (on bead) | 45 min | On bead, 1-2 hrs (Trypsin/LysC) | 1 (including SPE) | 1400 | 19.55 | 1.40 |
| Nanomics | 40 | Dilute (buffer), on bead incubation (60 min), washes (5 min) | 90 min | On bead, 1-2 hrs (Trypsin/LysC) | 1 | 2800 | 5.59 | 0.20 |
| PerCa | 50 | Precipitation, 20 for 15 min, centrifuge (60 min), HLB, lyophilise. SP3 digest | 120 min | ON, processed with Strap | 2 | 3500 | 4.17 | 0.12 |

*25µL of the total 100µL plasma lysis solution was taken from the original step (i.e., 25% of the ~140µg of plasma protein from the 2 µL input)*

**Supplementary Table 2:** Identification summaries reported by Spectronaut DirectDIA. Includes protein groups (PG), peptides (Pep) and precursors (Pre), as well as digestion efficiency (%) for (Ai) human biofluids—discovery method; (Aii) rat plasma—discovery method; (Bi) human biofluids—throughput method; (Bii) rat plasma—throughput method.

A) (i)

| Human, 25cm | Fluid | PG | Pep | Pre | Digest % | Pep/PG | Pre/PG |
|---|---|---|---|---|---|---|---|
| SP3 | Plasma | 1042 | 8973 | 13556 | 29.97 | 8.64 | 13.06 |
| | Serum | 1067 | 9188 | 13606 | 28.63 | 8.62 | 12.76 |
| Strap | Plasma | 1145 | 8646 | 12231 | 18.00 | 7.55 | 10.69 |
| | Serum | 1139 | 8918 | 12698 | 21.60 | 7.85 | 11.18 |
| PerCa | Plasma | 1662 | 10035 | 12323 | 18.33 | 6.05 | 7.43 |
| | Serum | 1600 | 10488 | 13368 | 19.03 | 6.55 | 8.35 |
| EnrichIST | Plasma | 1540 | 10580 | 14832 | 17.73 | 6.88 | 9.64 |
| | Serum | 1497 | 9602 | 13284 | 14.80 | 6.41 | 8.88 |
| Nanomics | Plasma | 1372 | 11222 | 14963 | 23.40 | 8.18 | 10.90 |
| | Serum | 1035 | 8275 | 11357 | 24.03 | 7.99 | 10.96 |
| SAX | Plasma | 1636 | 14594 | 20656 | 26.17 | 8.92 | 12.63 |
| | Serum | 1719 | 15111 | 21230 | 25.93 | 8.79 | 12.35 |
| HILIC | Plasma | 1664 | 14278 | 13081 | 23.37 | 8.62 | 11.86 |
| | Serum | 1740 | 15124 | 20991 | 21.93 | 8.69 | 12.06 |
| SAX/HILIC | Plasma | 1648 | 14872 | 20779 | 22.70 | 9.03 | 12.61 |
| | Serum | 1726 | 14375 | 19937 | 22.17 | 8.33 | 11.55 |

(ii)

| Rat, 25cm | PG | Pep | Pre | Digest % | Pep/PG | Pre/PG |
|---|---|---|---|---|---|---|
| SP3 | 1492 | 12371 | 17491 | 24.3 | 8.3 | 11.7 |
| Strap | 1515 | 11717 | 16002 | 18.2 | 7.7 | 10.5 |
| PerCa | 1524 | 9089 | 11553 | 17.9 | 6.0 | 7.6 |
| EnrichIST | 2539 | 17508 | 22358 | 18.6 | 6.9 | 8.8 |
| Nanomics | 2050 | 14027 | 18291 | 21.0 | 6.9 | 8.9 |
| MN SAX | 2863 | 24571 | 32501 | 21.4 | 8.6 | 11.4 |
| MN HILIC | 3058 | 27395 | 35945 | 19.8 | 9.0 | 11.8 |
| MN SAX/HILIC | 3003 | 26574 | 35035 | 20.4 | 8.8 | 11.7 |

B) (i)

| Human, 15cm | Fluid | PG | Pep | Pre | Digest % | Pep/PG | Pre/PG |
|---|---|---|---|---|---|---|---|
| SP3 | Plasma | 772 | 7568 | 10718 | 30.53 | 9.80 | 13.88 |
| | Serum | 813 | 7887 | 11024 | 29.52 | 9.71 | 13.57 |
| Strap | Plasma | 921 | 7617 | 10144 | 18.33 | 8.27 | 11.02 |
| | Serum | 891 | 7817 | 10509 | 22.07 | 8.79 | 11.82 |
| PerCa | Plasma | 1155 | 8361 | 9639 | 19.37 | 7.23 | 8.35 |
| | Serum | 1124 | 8239 | 9748 | 19.23 | 7.32 | 8.66 |
| EnrichIST | Plasma | 1077 | 8510 | 11230 | 18.46 | 7.90 | 10.43 |
| | Serum | 1029 | 7941 | 10448 | 16.43 | 7.72 | 10.15 |
| Nanomics | Plasma | 984 | 8837 | 11280 | 24.17 | 8.98 | 11.47 |
| | Serum | 766 | 6730 | 8664 | 25.20 | 8.80 | 11.32 |
| MN SAX | Plasma | 1165 | 12175 | 16459 | 27.60 | 10.45 | 14.13 |
| | Serum | 1210 | 12529 | 16928 | 26.43 | 10.39 | 14.00 |
| MN HILIC | Plasma | 1136 | 11582 | 15199 | 22.97 | 10.20 | 13.39 |
| | Serum | 1243 | 12171 | 16055 | 22.30 | 9.79 | 12.92 |
| MN SAX/HILIC | Plasma | 1145 | 11990 | 15832 | 23.15 | 10.48 | 13.83 |
| | Serum | 1186 | 11844 | 15664 | 22.87 | 9.99 | 13.21 |

(ii)

| Rat, 15cm | PG | Pep | Pre | Digest % | Pep/PG | Pre/PG |
|---|---|---|---|---|---|---|
| SP3 | 1160 | 10104 | 13598 | 25.00 | 8.72 | 11.73 |
| Strap | 1245 | 9485 | 12155 | 16.13 | 7.62 | 9.77 |
| PerCa | 1211 | 7067 | 8401 | 17.27 | 5.84 | 6.94 |
| EnrichIST | 1992 | 13358 | 16306 | 18.37 | 6.71 | 8.19 |
| Nanomics | 1623 | 11501 | 14099 | 22.13 | 7.09 | 8.70 |
| MN SAX | 2390 | 20699 | 26305 | 21.97 | 8.66 | 11.01 |
| MN HILIC | 2422 | 21103 | 26465 | 19.73 | 8.72 | 10.93 |
| MN SAX/HILIC | 2374 | 20688 | 26061 | 20.20 | 8.72 | 10.98 |

**Supplementary Table 3:** The counts of proteins in each workflow of higher intensity and enriched (↑ baseline) and lower intensity and depleted (↓ baseline) based on its STrap intensities as a baseline. The counts for 'nondetect baseline' are for proteins that were not present in the STrap baseline, and enriched in that workflow. Tables in (A) include for (i) human biofluids—discovery method; (ii) rat plasma—discovery method; tables in (B) (ii) human biofluids—throughput method; (iI) rat plasma—throughput method.

A) (i)

| Human, 25cm | Fluid | ↑ baseline | ↓ baseline | nondetect baseline |
|---|---|---|---|---|
| EnrichIST | Plasma | 547 | 377 | 580 |
| | Serum | 592 | 315 | 587 |
| HILIC | Plasma | 375 | 589 | 734 |
| | Serum | 432 | 538 | 780 |
| SAX | Plasma | 426 | 516 | 686 |
| | Serum | 398 | 601 | 738 |
| SAX/HILIC | Plasma | 407 | 510 | 730 |
| | Serum | 411 | 564 | 740 |
| Nanomics | Plasma | 397 | 454 | 525 |
| | Serum | 330 | 376 | 309 |
| PerCa | Plasma | 675 | 281 | 745 |
| | Serum | 646 | 366 | 588 |
| SP3 | Plasma | 213 | 709 | 97 |
| | Serum | 272 | 665 | 115 |

(ii)

| Rat, 25cm | Fluid | ↑ baseline | ↓ baseline | nondetect baseline |
|---|---|---|---|---|
| EnrichIST | Plasma | 834 | 450 | 1265 |
| HILIC | Plasma | 684 | 703 | 1735 |
| SAX | Plasma | 637 | 724 | 1557 |
| SAX/HILIC | Plasma | 662 | 724 | 1679 |
| Nanomics | Plasma | 581 | 733 | 770 |
| PerCa | Plasma | 758 | 275 | 436 |
| SP3 | Plasma | 358 | 893 | 159 |

B) (i)

| Human, 15cm | Fluid | ↑ baseline | ↓ baseline | nondetect baseline |
|---|---|---|---|---|
| EnrichIST | Plasma | 430 | 341 | 340 |
| | Serum | 448 | 307 | 316 |
| HILIC | Plasma | 363 | 427 | 398 |
| | Serum | 363 | 473 | 488 |
| SAX | Plasma | 342 | 493 | 393 |
| | Serum | 327 | 502 | 454 |
| SAX/HILIC | Plasma | 351 | 442 | 410 |
| | Serum | 349 | 459 | 451 |
| Nanomics | Plasma | 737 | 344 | -56 |
| | Serum | 258 | 351 | 174 |
| PerCa | Plasma | 544 | 289 | 387 |
| | Serum | 511 | 330 | 353 |
| SP3 | Plasma | 161 | 595 | 47 |
| | Serum | 183 | 597 | 61 |

(ii)

| Rat, 15cm | Fluid | ↑ baseline | ↓ baseline | nondetect baseline |
|---|---|---|---|---|
| EnrichIST | Plasma | 676 | 371 | 945 |
| HILIC | Plasma | 574 | 565 | 1324 |
| SAX | Plasma | 550 | 593 | 1291 |
| SAX/HILIC | Plasma | 552 | 601 | 1284 |
| Nanomics | Plasma | 489 | 550 | 583 |
| PerCa | Plasma | 613 | 231 | 326 |
| SP3 | Plasma | 191 | 832 | 108 |

### Supplementary Figures

#### Supplementary Figure 1:

A)

| Dataset | Precursors | Peptides | Proteins | Pep/PG | Pre/PG |
|---|---|---|---|---|---|
| E01 – Neat | 10,335 | 7,094 | 602 | 11.6 | 17.0 |
| E01 – PerCa | 6,235 | 4,798 | 837 | 5.47 | 7.01 |
| E01 – MN SAX | 14,247 | 11,130 | 1202 | 9.09 | 9.63 |
| E05 – Neat | 20,732 | 14,289 | 1221 | 9.61 | 13.9 |
| E05 – PerCa | 15,326 | 12,091 | 2186 | 4.12 | 5.08 |
| E05 – MN SAX | 69,332 | 53,887 | 4559 | 7.71 | 9.63 |

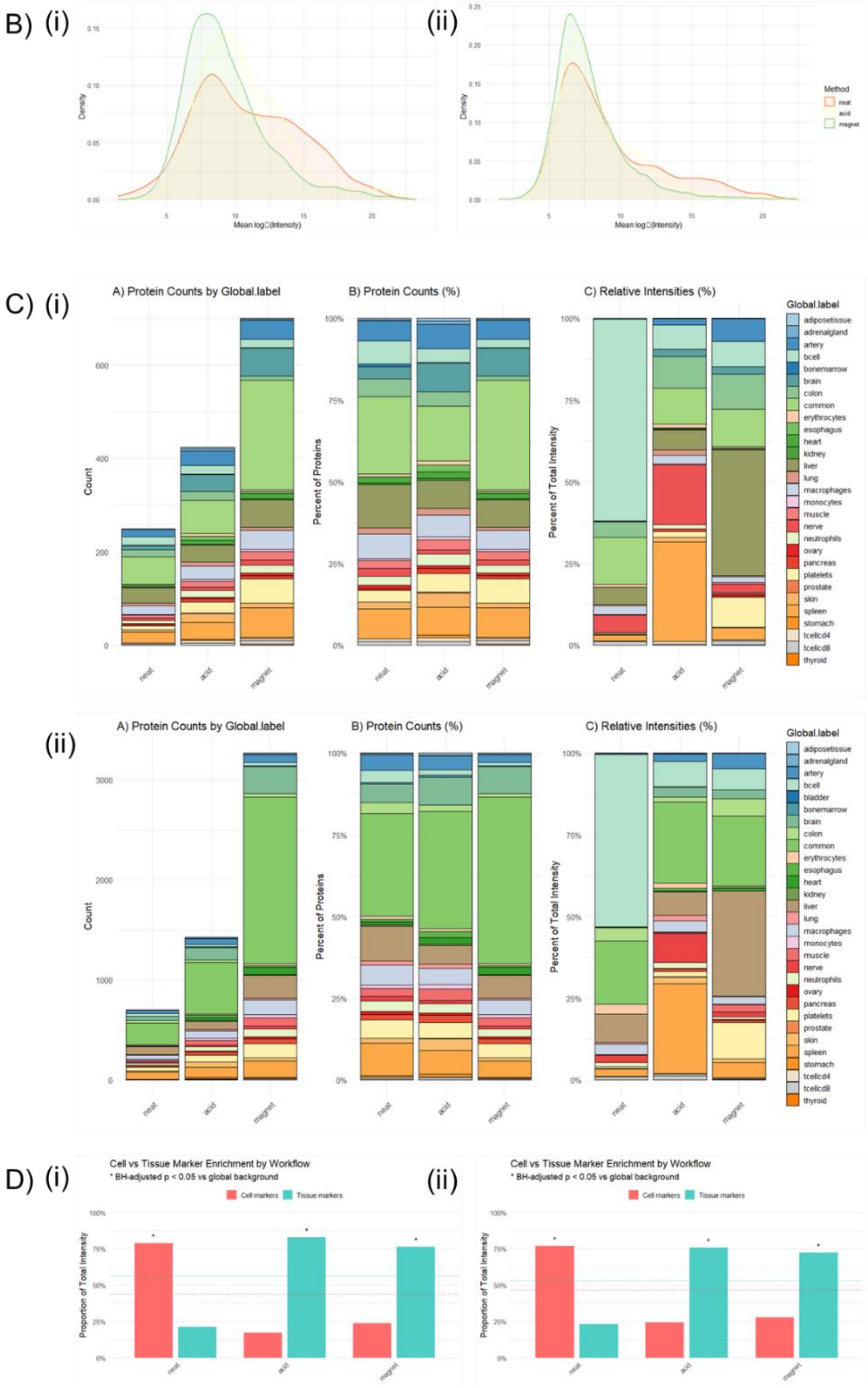

**Supplementary Figure 1.** Inter-benchmarking comparisons between shared workflows in this study and Beimers et al (1), specifically Neat, MagNet SAX and PerCa. Data was downloaded from PRIDE, and the precursor-level Spectronaut data for the three workflows was processed for Experiment 1 (E01; technical control, plasma pools) and Experiment 5 (E05) for the non-small cell lung cancer (NSCLC) cohort. (A) Table of identification summaries for comparison with Supplementary Table 2. (B) Kernel density curves for (i) E01 and (ii) E02). (C) Stacked bar charts summarising for each workflow (left to right) raw protein counts per global label, percent of protein counts per label, and percent of total intensity per label in (i) E01 and (ii) E05. (C) Cell- versus tissue-marker enrichment by workflow for (i) E01 (ii) E05. Grouped bars show the proportion of total marker intensity assigned to cell markers and to tissue markers for each method. Dashed horizontal lines indicate expected background proportions estimated from pooled intensities across all samples. Enrichment is assessed with one-sided exact binomial tests (rounded summed intensities treated as counts), with Benjamini–Hochberg adjustment across methods; asterisks denote adjusted $p < 0.05$ for either cell- or tissue-marker enrichment.

**Supplementary Figure 2:**

A)

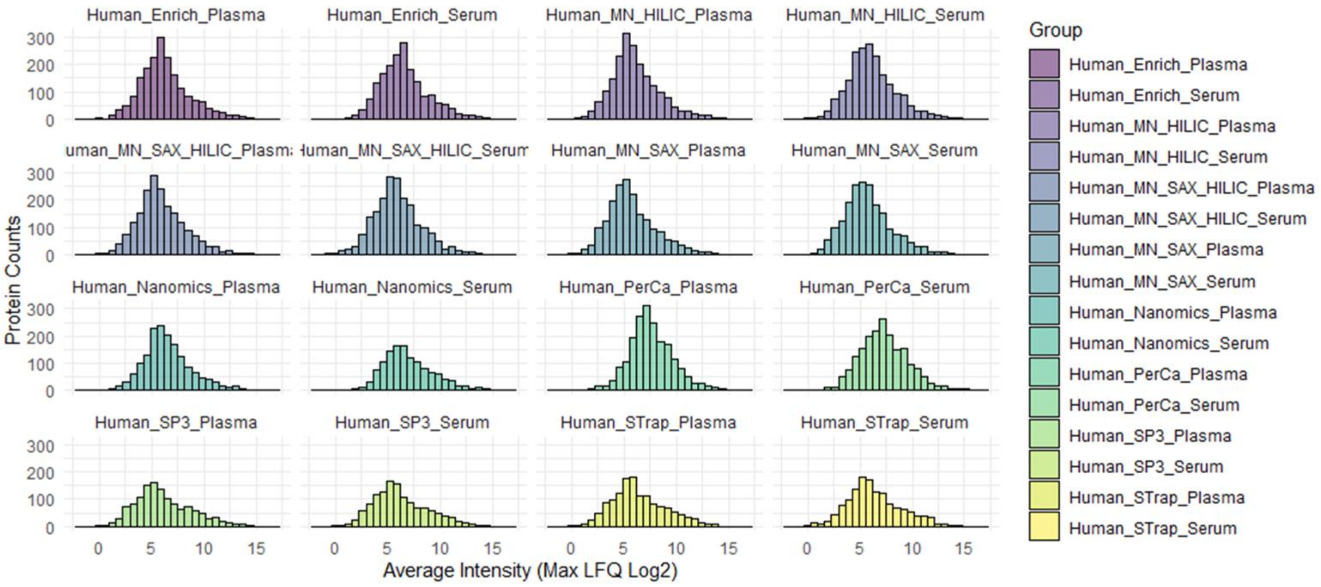


B)

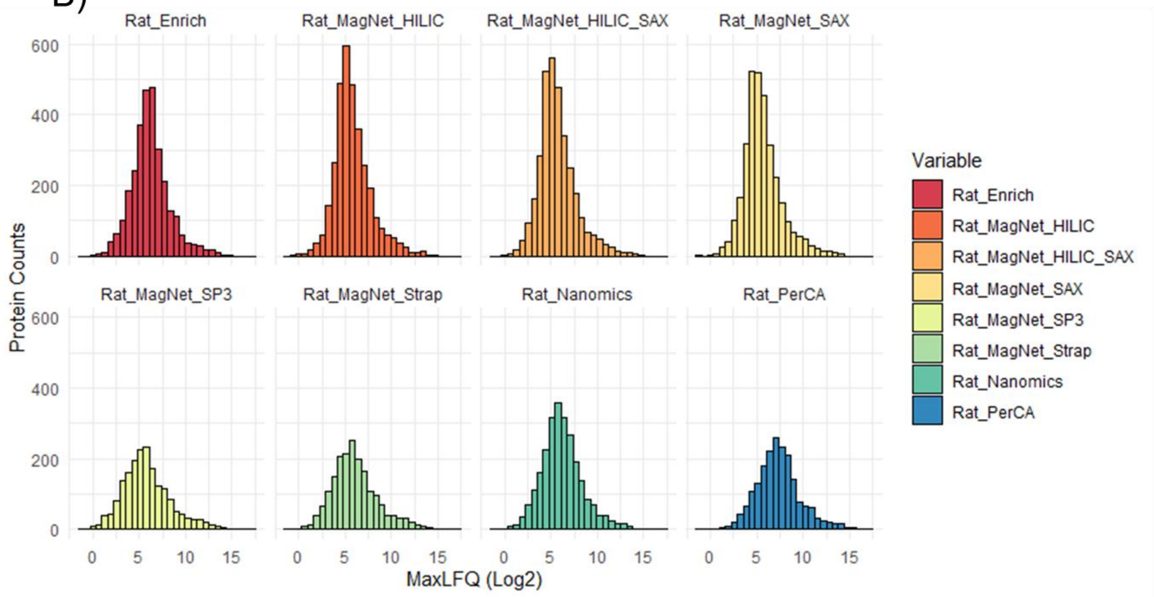

**Supplementary Figure 2.** Binned histograms (protein counts per intensity bin) faceted separately for workflow in the **discovery method** in human biofluids (A) and rat plasma (B).

**Supplementary Figure 3:**

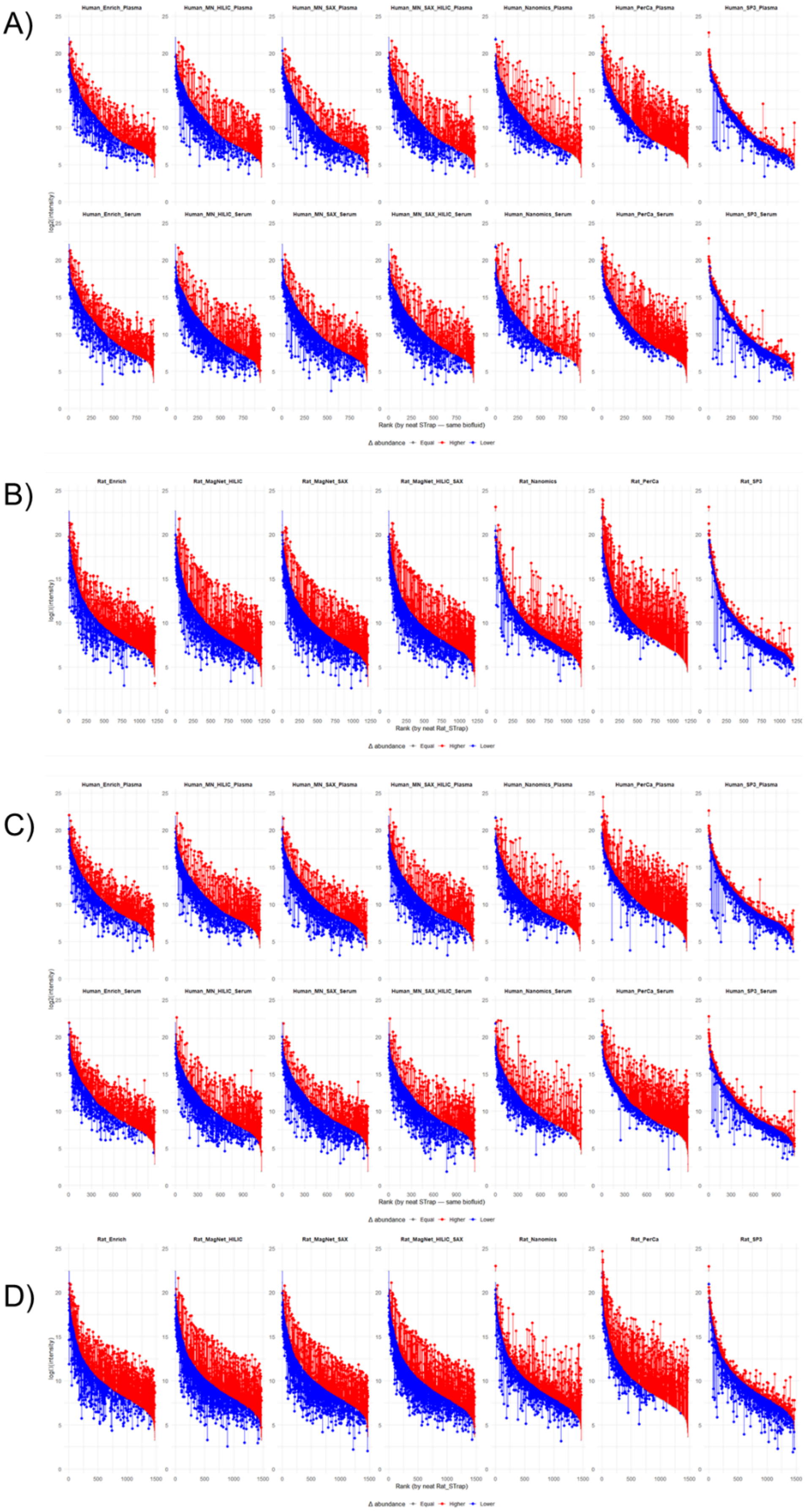

**Supplementary Figure 3.** Rank-abundance plots of all proteins detected in baselines as defined by the average log2 replicate intensities from STrap. For each protein, the average log2 intensity in each workflow is plotted as a point; a segment connects the STrap baseline to the workflow value, coloured red if higher than baseline and blue if lower. Plots are shown for (A) human biofluids—discovery method; (B) rat plasma—discovery method; tables in (C) human biofluids—throughput method; (D) rat plasma—throughput method

#### Supplementary Figure 4:

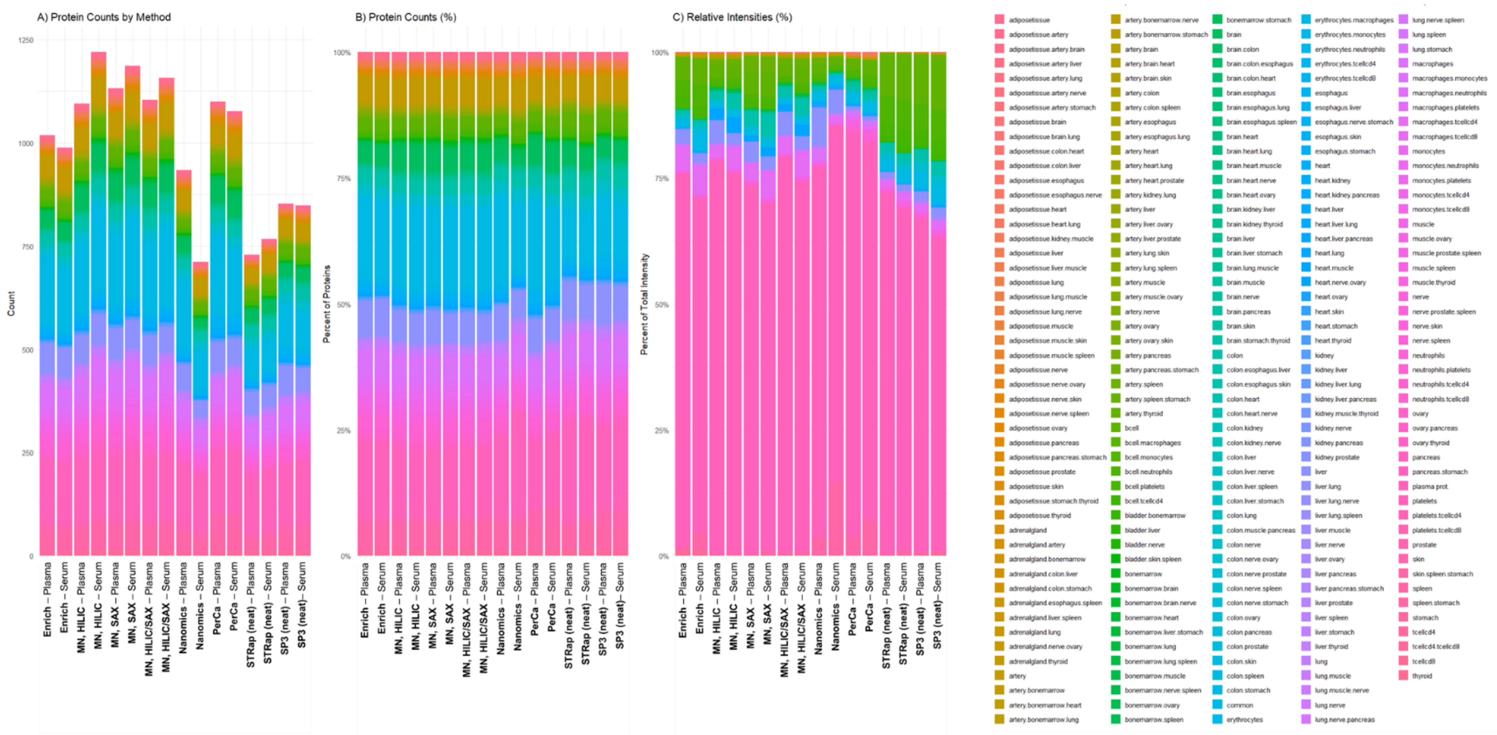

**Supplementary Figure 4.** Stacked bar charts summarising for each workflow, (A) raw protein counts per global label, (B) percent of protein counts per label, and (C) percent of total intensity across all global.label categories in the atlas (2), irrespective of the combination/total tissue and cell maps the protein belongs to. Intensities are summed within method × label after back-transformation from log2 and expressed as a proportion of each method's total intensity (0–100% scale).

**Supplementary Figure 5:**

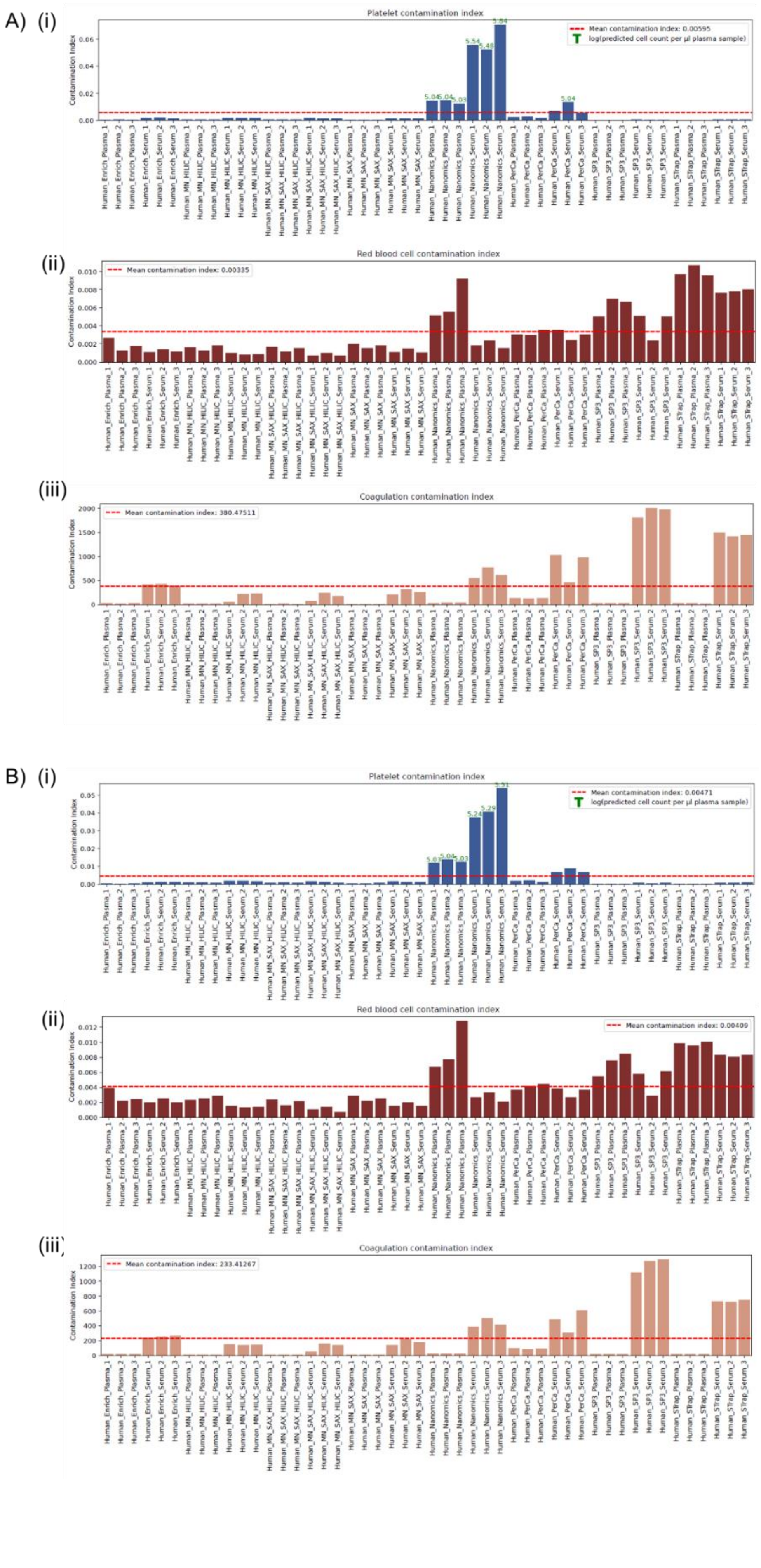

**Supplementary Figure 5.** Evaluation of samples for platelet contamination, erythrocyte lysis, and residual coagulation protein carryover using Baize v3.0 in the Throughput data (A) and Discovery data (B). Calculation of cell-type specific contamination indices is described in Guo et al (3). Here, platelet contamination was detected in ProteoNano plasma and serum samples in the 15cm data (A(i)) alongside in one PerCa serum replicate, and ProteoNano samples in the 25cm data (panel B(i)). No RBC contamination was detected (A(ii), B(ii). In the coagulation panel, higher contamination indexes were seen in all serum samples, as expected, with higher indexes in Neat workflows, followed by PerCa and ProteoNano, while EnrichIST and MagNet workflows were all below the average in the datasets.

#### Supplementary Figure 6:

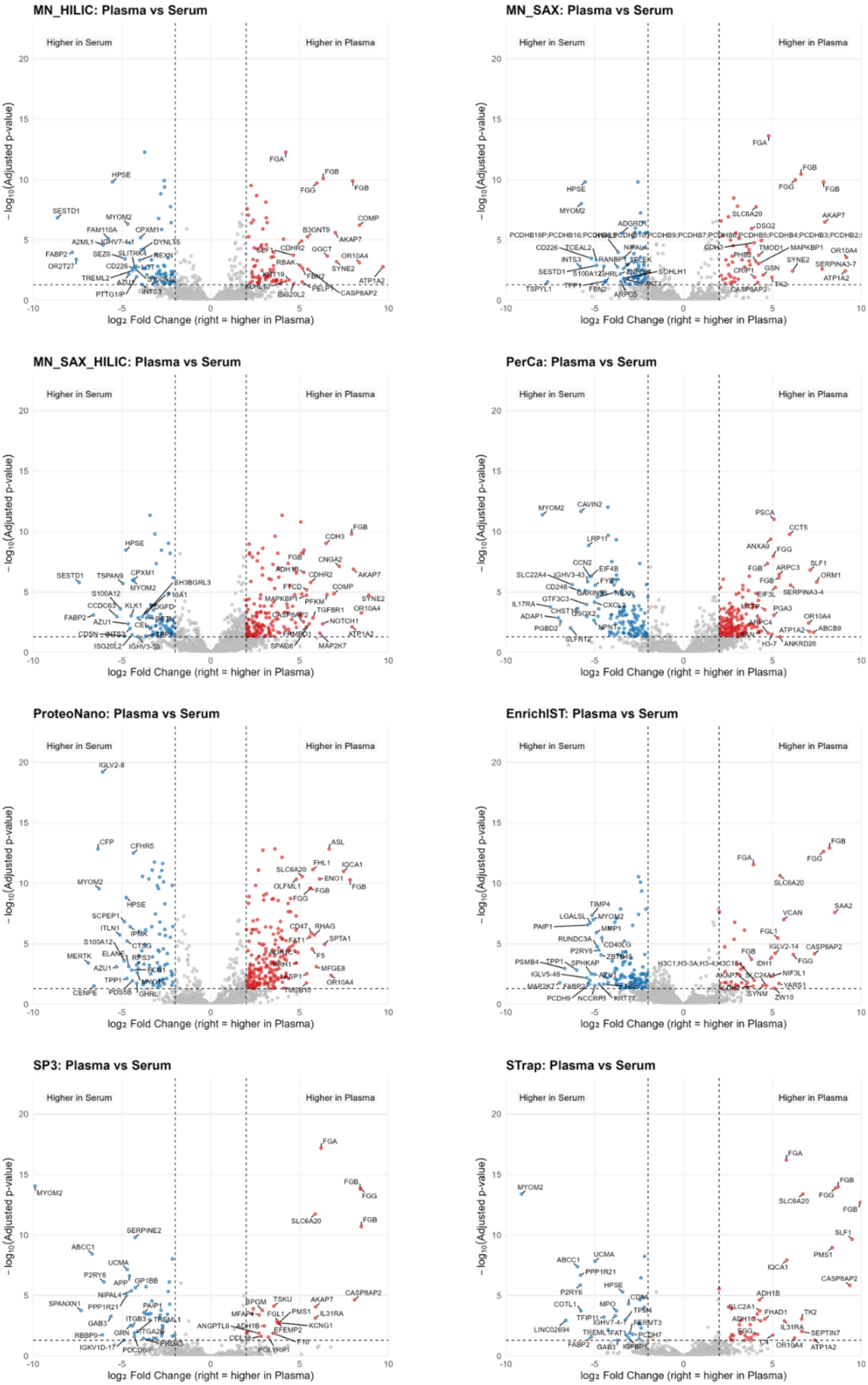

**Supplementary Figure 6.** Volcano plots for inter-biofluid pairwise comparisons in each workflow (i.e., plasma vs serum). Axes show $\log_2$ fold change (x) versus $-\log_{10}$(adjusted p) (y). Dashed vertical lines indicate fold-change thresholds ($\log_2FC = 2$) and the dashed horizontal line indicates the significance threshold (FDR-adjusted p = 0.05). Points are coloured by significance category: higher in serum (blue), higher in the plasma (red), and not significant (grey). The 20 most extreme significant proteins on each side (largest $\log_2FC$, then lowest p) are labelled by gene name.

#### Supplementary Figure 7:

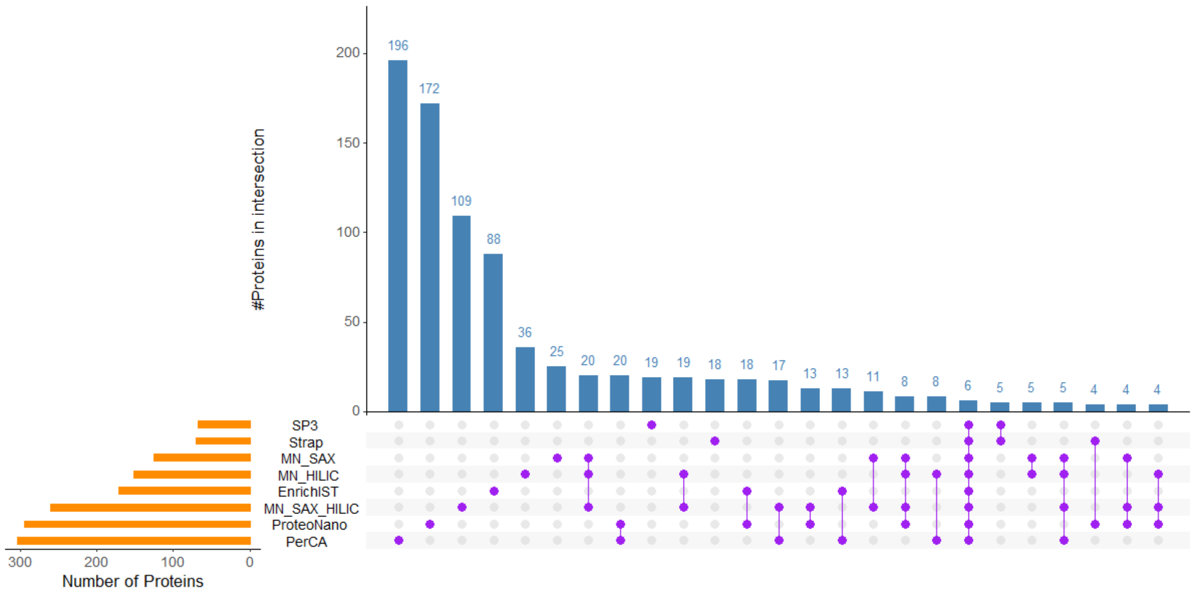

**Supplementary Figure 7.** UpSet plot of statistically significant differentially abundant proteins in plasma vs serum pairwise tests within workflows. upSET is arranged by frequency, and the top 25 intersections are shown.

#### Supplementary Figure 8:

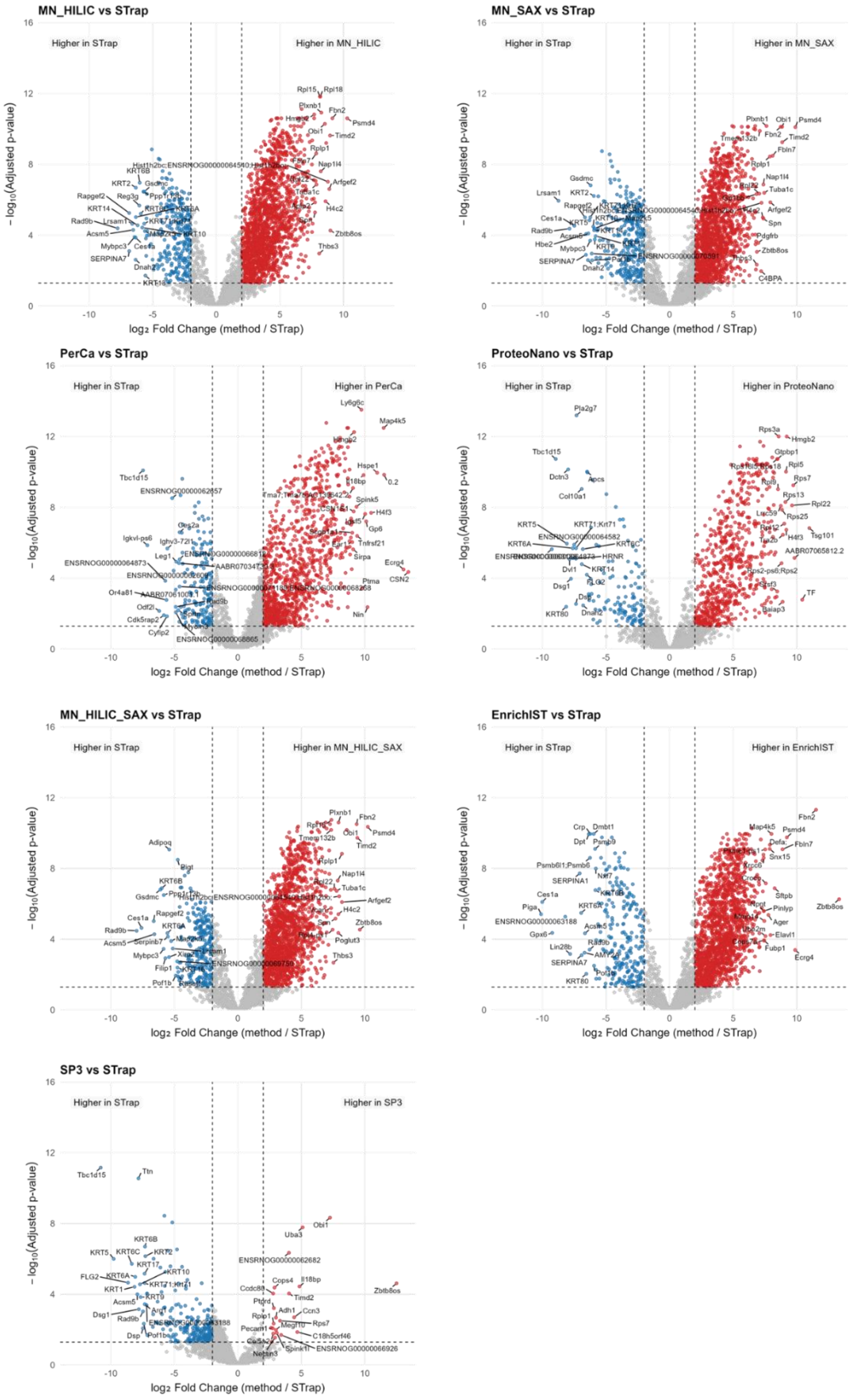

**Supplementary Figure 8.** Volcano plots for pairwise comparisons in rat plasma of each workflow to the STrap baseline. Axes show $\log_2$ fold change (x) versus $-\log_{10}$(adjusted p) (y). Dashed vertical lines indicate fold-change thresholds ($\log_2 FC = 2$) and the dashed horizontal line indicates the significance threshold (FDR-adjusted p = 0.05). Points are coloured by significance category: higher in serum (blue), higher in the plasma (red), and not significant (grey). The 20 most extreme significant proteins on each side (largest $\log_2 FC$, then lowest p) are labelled by gene name.

**Supplementary Figure 9:**

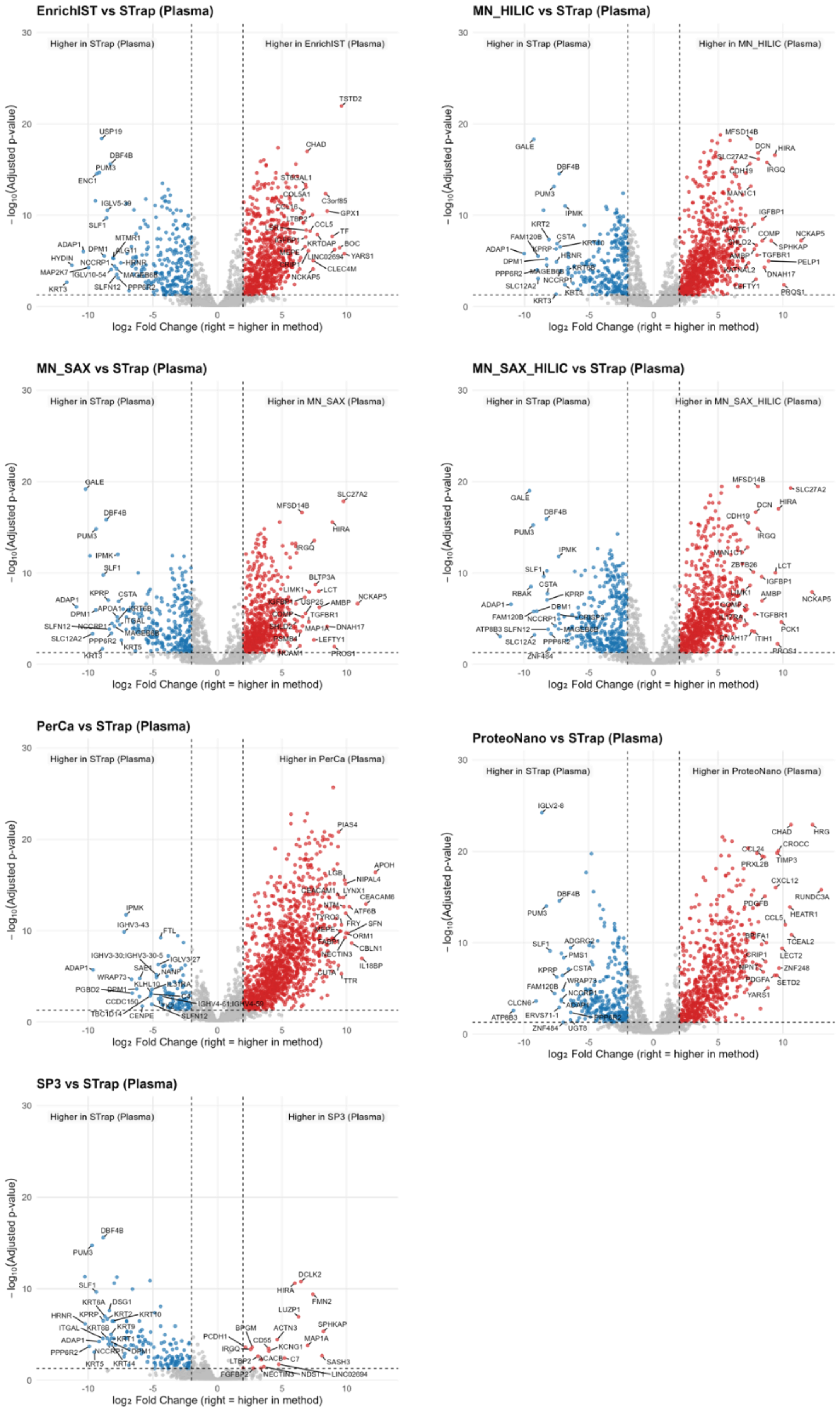

**Supplementary Figure 9.** Volcano plots for pairwise comparisons in human plasma of each workflow to the STrap baseline. Axes show $\log_2$ fold change (x) versus $-\log_{10}$(adjusted p) (y). Dashed vertical lines indicate fold-change thresholds ($\log_2$FC = 2) and the dashed horizontal line indicates the significance threshold (FDR-adjusted p = 0.05). Points are coloured by significance category: higher in serum (blue), higher in the plasma (red), and not significant (grey). The 20 most extreme significant proteins on each side (largest $\log_2$FC, then lowest p) are labelled by gene name.

**Supplementary Figure 10:**

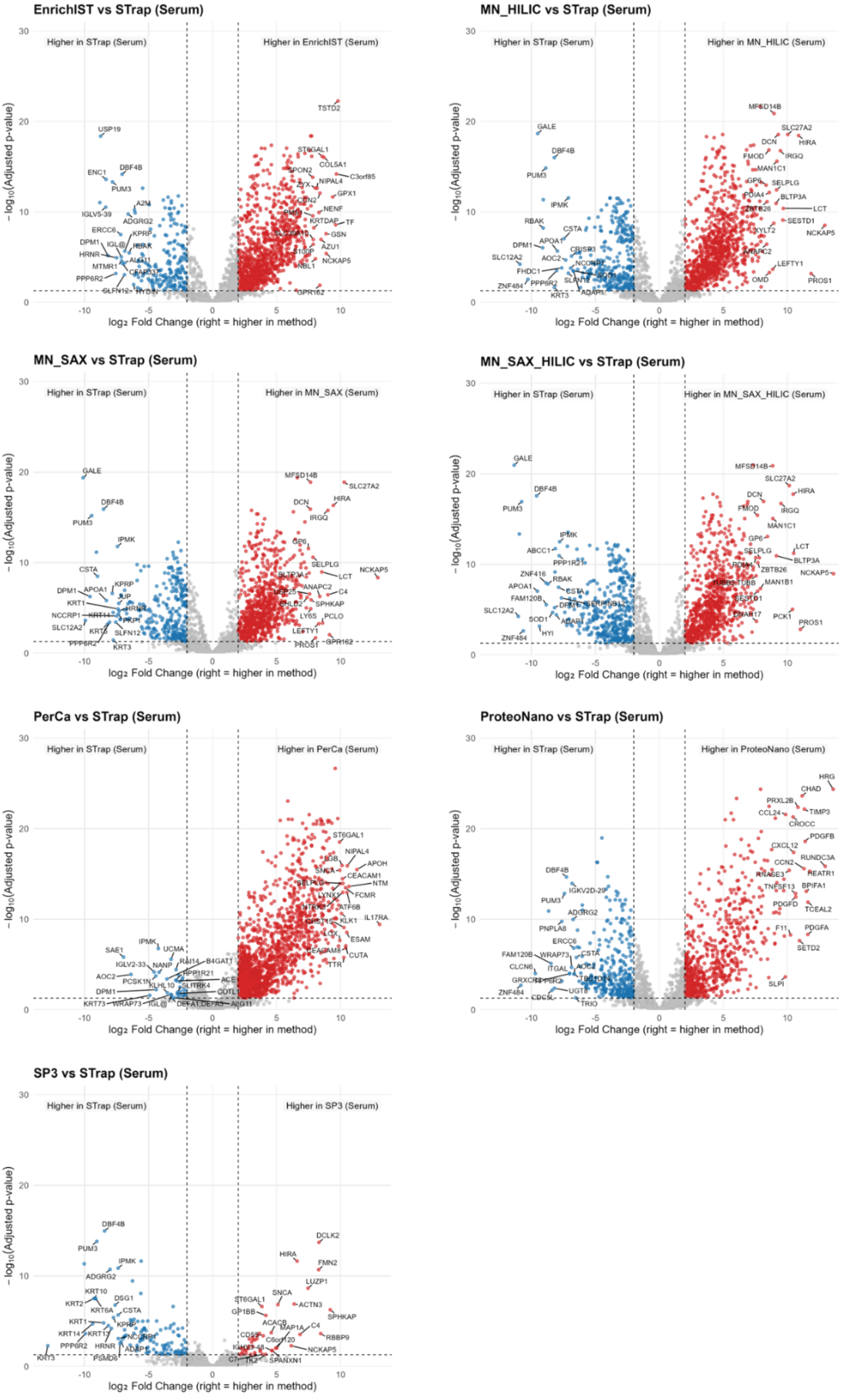

**Supplementary Figure 10.** Volcano plots for pairwise comparisons in human serum of each workflow to the STrap baseline. Axes show $\log_2$ fold change (x) versus $-\log_{10}$(adjusted p) (y). Dashed vertical lines indicate fold-change thresholds ($\log_2$FC = 2) and the dashed horizontal line indicates the significance threshold (FDR-adjusted p = 0.05). Points are coloured by significance category: higher in serum (blue), higher in the plasma (red), and not significant (grey). The 20 most extreme significant proteins on each side (largest $\log_2$FC, then lowest p) are labelled by gene name.

**Supplementary Figure 11:**

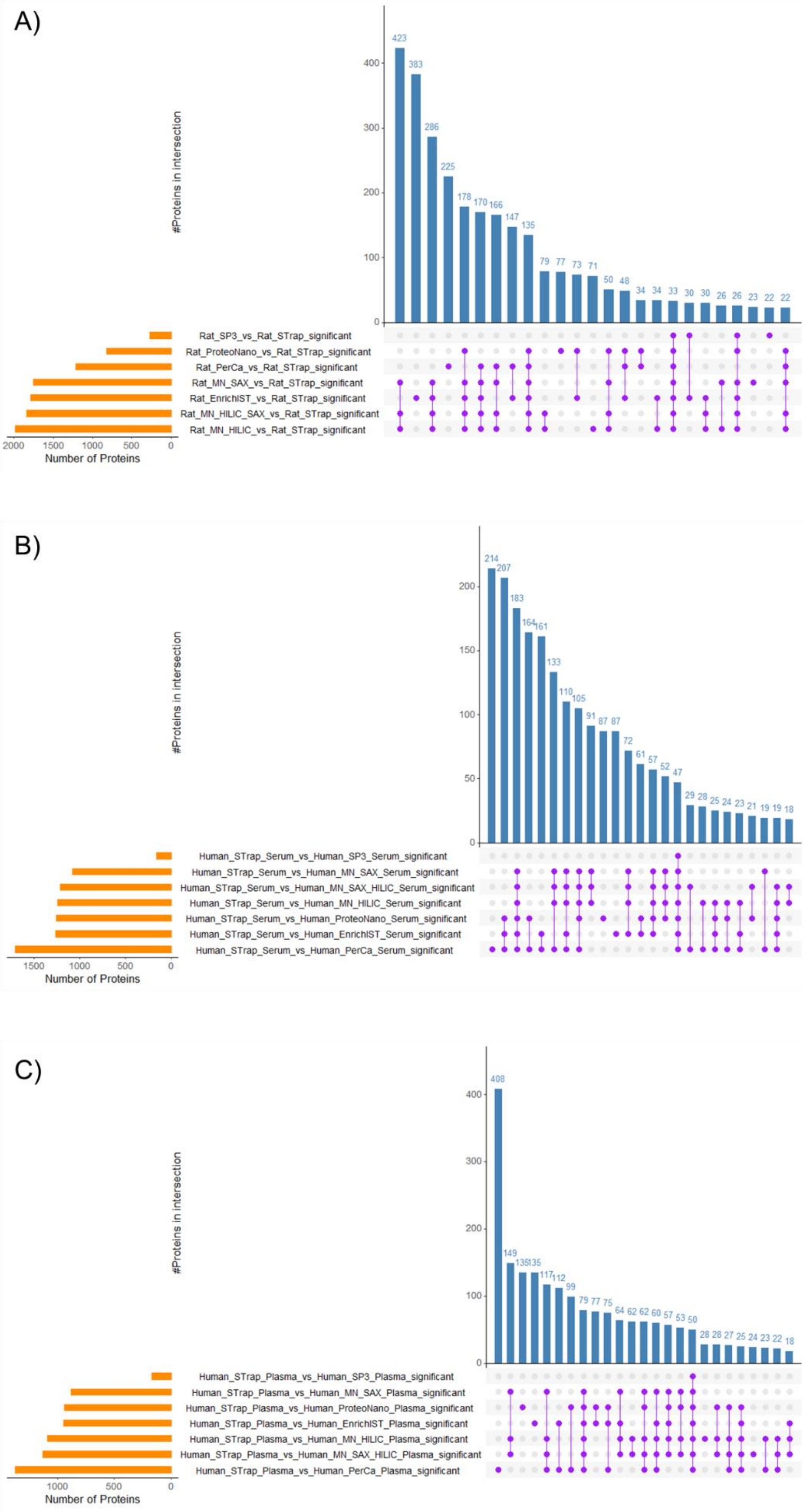

**Supplementary Figure 11.** upSET plot of statistically significant differentially abundant proteins in workflow vs Strap baseline pairwise tests in the **discovery method** for (A) rat plasma, (B) human serum, and (C) human plasma. upSET is arranged by frequency, and the top 25 intersections are shown.

**Supplementary Figure 12:**

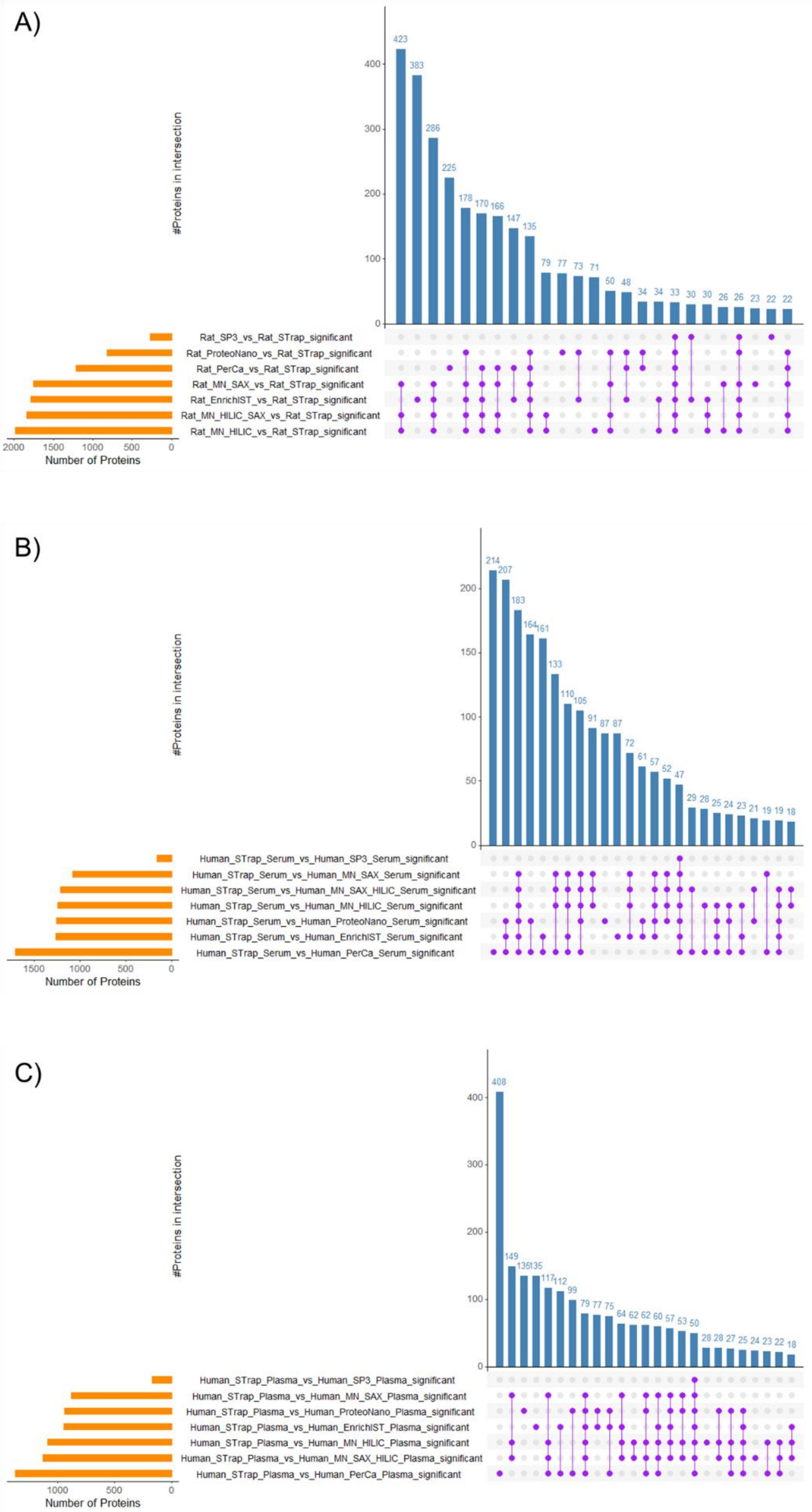

**Supplementary Figure 12.** upSET plot of statistically significant differentially abundant proteins in workflow vs Strap baseline pairwise tests in the **throughput method** for (A) rat plasma, (B) human serum, and (C) human plasma. upSET is arranged by frequency, and the top 25 intersections are shown.

### Supplementary Figure 13:

#### A) *Rat, 25cm, Discovery, Figure 6A, Cluster 1*

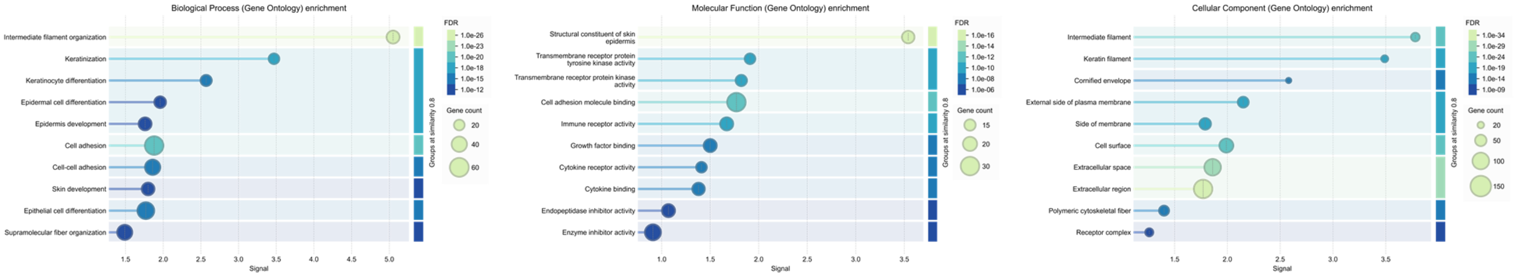


#### B) *Rat, 25cm, Discovery, Figure 6A, Cluster 2*

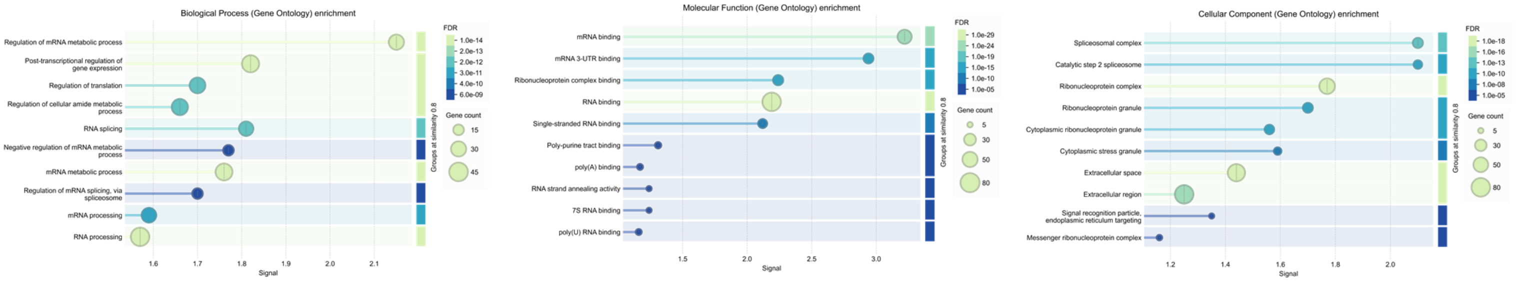


#### C) *Rat, 25cm, Discovery, Figure 6A, Cluster 3*

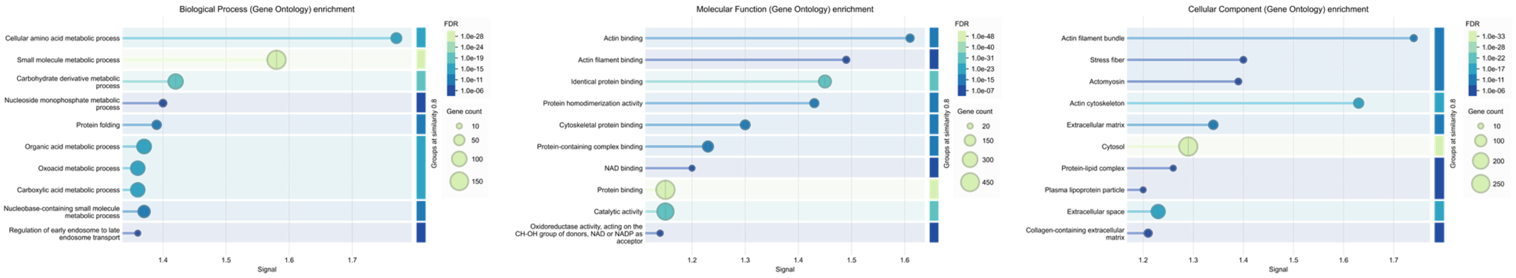


#### D) *Rat, 25cm, Discovery, Figure 6A, Cluster 4*

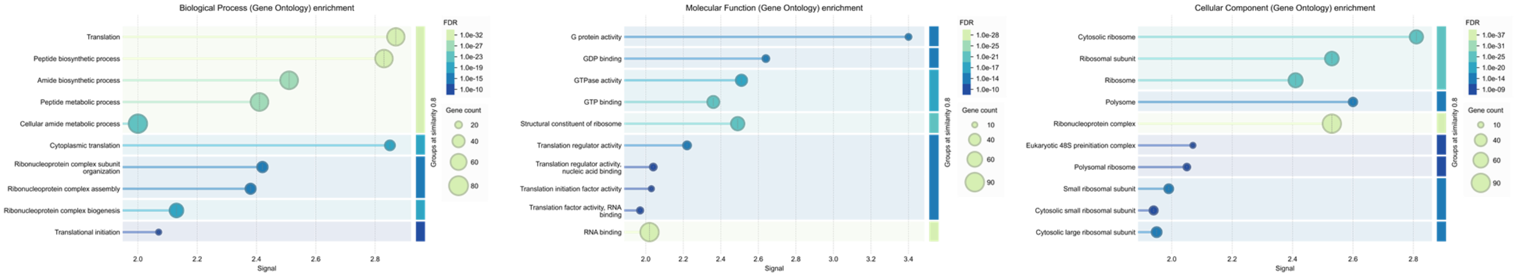


#### E) *Rat, 25cm, Discovery, Figure 6A, Cluster 5*

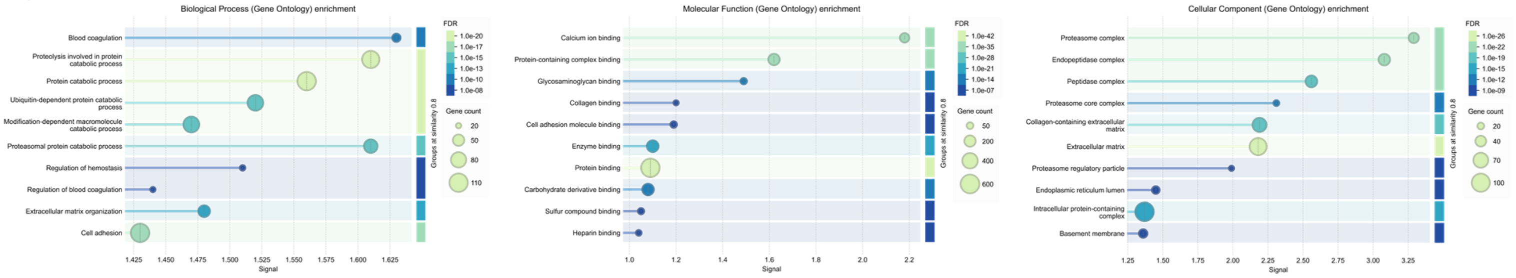

**Supplementary Figure 13.** STRING DB functional enrichment across clusters of interest based on the hierarchical clustering by DIA Analyst. Bubble heatmaps summarising top 10 terms based on (left – right: GO:BP, GO:MF, GO:CC). Circle area is proportional to the number of mapped proteins (gene count), and fill colour encodes the FDR-corrected enrichment q-value from STRING (darker = stronger enrichment). Vertical gridlines delineate workflows. All results are from STRING functional enrichment with multiple-testing correction (FDR); only the top terms per workflow are shown. Panel titles from (A) – (G) indicate which cluster from rat were submitted (Figure 6A).

**Supplementary Figure 14:**

A)

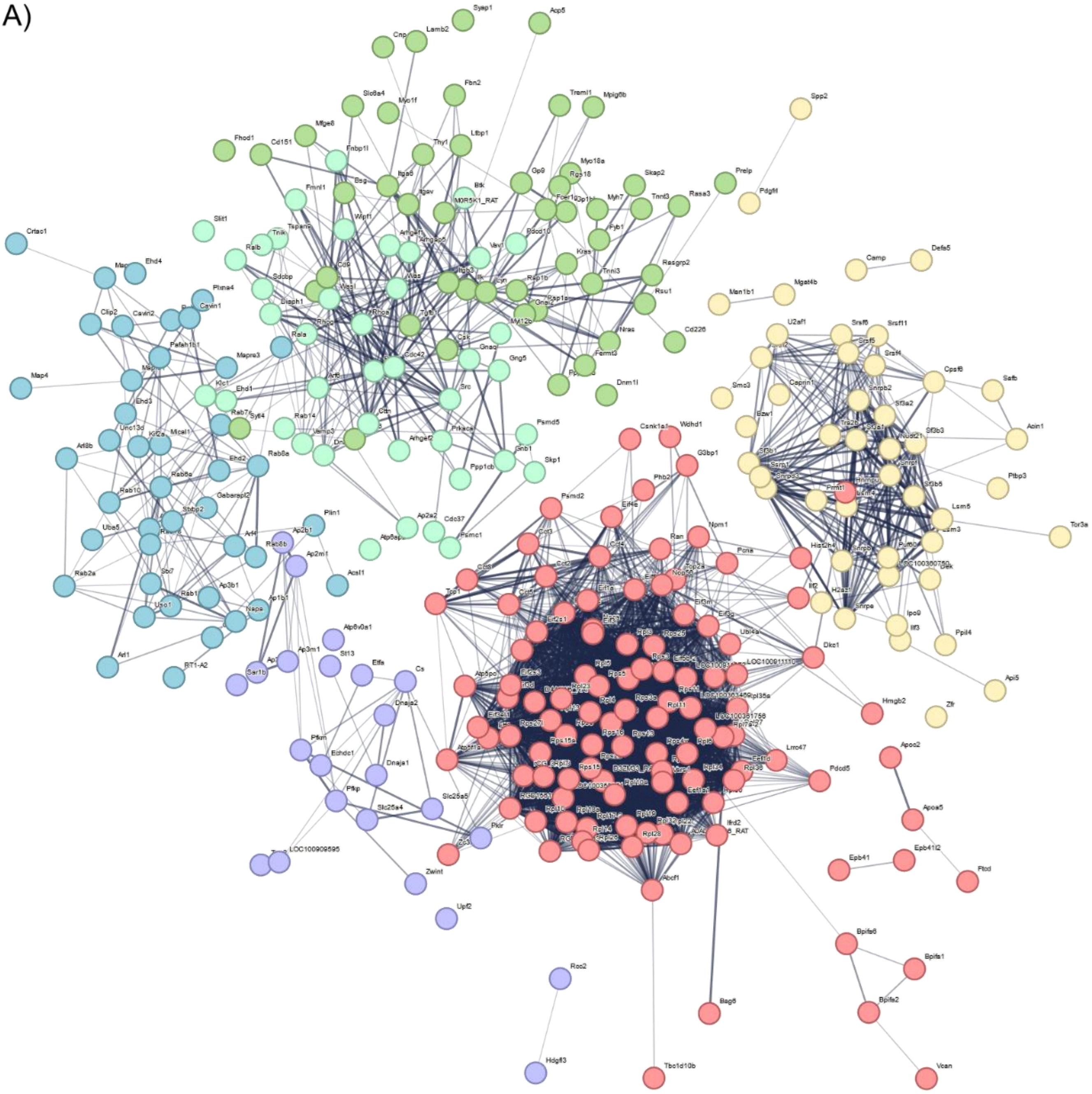

B)

| color | cluster Id | gene count | description |
|---|---|---|---|
| | Cluster 1 | 116 | GTP hydrolysis and joining of the 60S ribosomal subunit |
| | Cluster 2 | 61 | Spliceosome |
| | Cluster 3 | 53 | Platelet activation, signaling and aggregation |
| | Cluster 4 | 42 | - 1. Yersinia infection<br>2. Actin filament organization, and RAC1 GTPase cycle |
| | Cluster 5 | 42 | - 1. Endosomal transport<br>2. GTP-binding |
| | Cluster 6 | 39 | 4933400A11Rik, Ap2b1, Ap2m1, Ap3d1, Ap3m1, Atp6v0a1, Cs, Dnaja1, Dnaja2, Ech... |

**Supplementary Figure 14.** STRING-DB network of proteins in Rat Plasma cluster 4 (Figure 6A) depicting functional relationships. Settings include K-means clustering set for 6 clusters, with dotted lines indicating the edges between clusters. All disconnected nodes have been hidden. Below, the colour of the five clusters, gene count and a description of the grouped gene functions.

#### Supplementary Figure 15:

A) *Human, 25cm, Discovery, Figure 7A, Cluster 1*

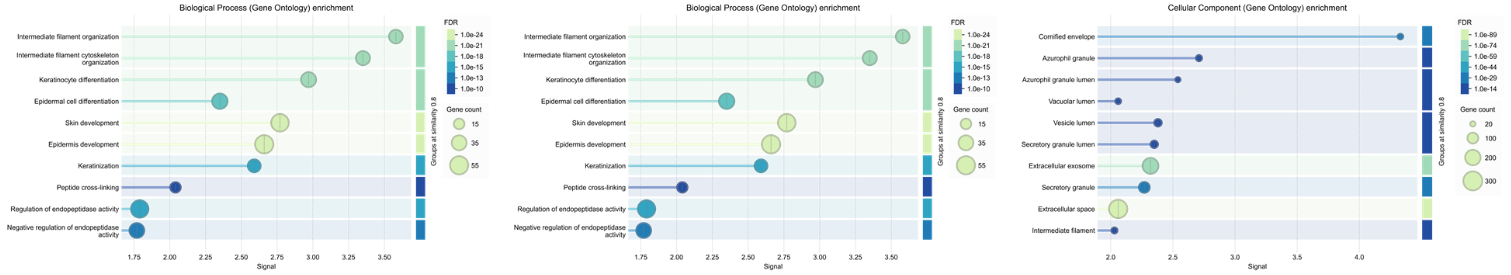


B) *Human, 25cm, Discovery, Figure 7A, Cluster 5*

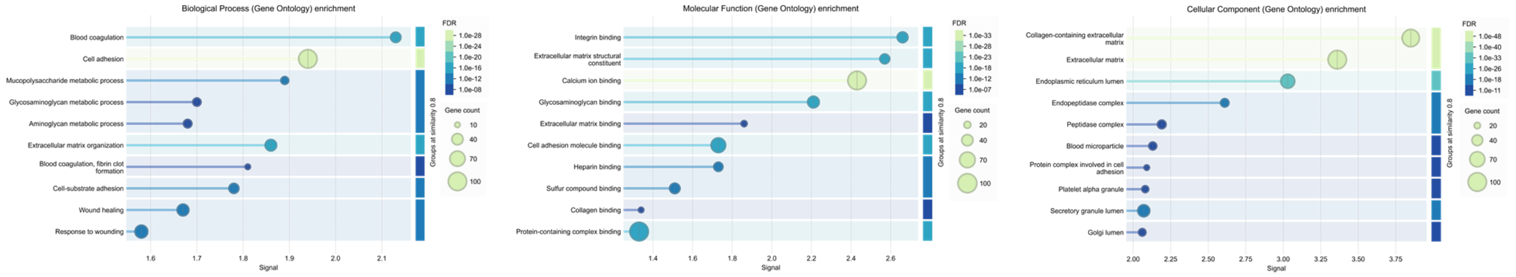

**Supplementary Figure 15:** STRING DB functional enrichment across clusters of interest based on the hierarchical clustering by DIA Analyst. Bubble heatmaps summarising top 10 terms based on (left – right: GO:BP, GO:MF, GO:CC). Circle area is proportional to the number of mapped proteins (gene count), and fill colour encodes the FDR-corrected enrichment q-value from STRING (darker = stronger enrichment). Vertical gridlines delineate workflows. All results are from STRING functional enrichment with multiple-testing correction (FDR); only the top terms per workflow are shown. Panel titles from (A) – (C) indicate which cluster from human were submitted (Figure 7A)
